# Structure-based Design of CDC42 Effector Interaction Inhibitors For the Treatment of Cancer

**DOI:** 10.1101/2021.10.14.464305

**Authors:** Sohail Jahid, Jose A. Ortega, Linh M. Vuong, Isabella Maria Acquistapace, Stephanie J. Hachey, Jessica L. Flesher, Maria Antonietta La Serra, Nicoletta Brindani, Giuseppina La Sala, Jacopo Manigrasso, Jose M. Arencibia, Sine Mandrup Bertozzi, Maria Summa, Rosalia Bertorelli, Andrea Armirotti, Rongsheng Jin, Zheng Liu, Chi-Fen Chen, Robert Edwards, Christopher C.W. Hughes, Marco De Vivo, Anand K. Ganesan

**Author notes:** These authors contributed equally to experimental work (Sohail Jahid, Jose A. Ortega, Linh M. Vuong). These collaborating senior authors equally contributed to study design, data analysis, and manuscript preparation (Marco De Vivo, Anand K. Ganesan).

## Abstract

CDC42 family GTPases (RHOJ, RHOQ, CDC42) are upregulated but rarely mutated in cancer and control both the ability of tumor cells to invade surrounding tissues and the ability of endothelial cells to vascularize tumors. Here we use computer-aided drug design to discover a new chemical entity (ARN22089) that targets CDC42 GTPases and blocks CDC42 effector interactions without affecting the binding between closely related GTPases (RAC1, RAS, RAL) and their downstream effectors. Our lead compound has broad activity against a panel of cancer cell lines, inhibits S6 phosphorylation and MAPK activation, activates pro-inflammatory and apoptotic signaling, and blocks tumor growth and angiogenesis in three-dimensional vascularized microtumor models (VMT) *in vitro*. In addition, ARN22089 has a favorable pharmacokinetic profile and can inhibit the growth of BRAF mutant mouse melanomas and patient-derived xenografts *in vivo*. Taken together, this work identifies a promising new class of therapeutic agents that influence tumor growth by modulating CDC42 signaling in both the tumor cell and its microenvironment.

## INTRODUCTION

CDC42 GTPases (RHOJ, RHOQ, CDC42) are linked to multiple human cancers and modulate cell-cycle progression, tumor cell migration/invasion, and tumor angiogenesis.^1^ RHOJ, a known regulator of melanoma^2^, breast cancer^3^, and gastric cancer^4^ progression, activates signaling cascades in endothelial cells that are required for tumor angiogenesis^5–7^ and induces PAK signaling in tumor cells to promote growth.^2^ Similarly, CDC42 is a critical regulator of angiogenic sprouting^8^ and tubulogenesis^9^ in endothelial cells and promotes tumor cell proliferation and migration.^10^ In addition, CDC42 and its downstream PAK kinases are regulators of MAPK inhibitor resistance in BRAF mutant melanoma.^11^ Finally, RHOQ, while less studied, is also known to promote tumor invasion^12^ and angiogenesis^13^.

The GTPase activity of CDC42 family members is tightly regulated by *i)* guanine nucleotide exchange factors (GEFs); *ii)* guanine nucleotide dissociation inhibitors (GDIs); and *iii)* GTPase activating proteins (GAPs).^14^ CDC42 activity is also controlled by post-translational modification, including prenylation, which controls membrane localization.^14^ CDC42 family members are considered “undruggable” due to their globular structure with limited small molecule binding pockets, and their high affinity for GTP/GDP.^1^ Moreover, the tight regulation of CDC42 GTPases by GEFs, GAPs, and GDIs adds an additional level of complexity to targeting these proteins. Nonetheless, recent efforts have focused on developing small molecules that inhibit CDC42 signaling. Existing inhibitors block the ability of GEFs to activate RAC/CDC42^15^, prevent RAC/CDC42 from localizing to membranes^16^, or block RAC/CDC42 GTP binding^17^. Unfortunately, the efficacy of these strategies is limited by the poor selectivity and/or poor bioavailability of the inhibitors.^1^ Furthermore, as CDC42 and RAC1 have overlapping roles in platelet function^18^, targeting the signaling of both of these GTPases simultaneously often results in thrombocytopenia *in vivo*^19^. RAC1 also plays an important role in the cardiovascular system, and targeting RAC1 results in cardiotoxicity.^20^ New structure-based targeting strategies are needed to develop selective drug-like inhibitors that target the CDC42 family without targeting RAC1 to eliminate potential side effects.

Recent pioneering work has identified structural pockets in G12C mutant RAS and developed inhibitors that can covalently modify and inhibit G12C RAS function.^21,22^ While these approaches can be applied to RAS mutant cancers, they cannot be adapted to target CDC42 as these proteins are rarely mutated. Here we focus on CDC42 and RHOJ, two CDC42 family members that have well-defined roles in cancer. Using a CDC42-PAK6 crystal structure (PDBid: 2ODB), we generate a RHOJ-PAK1 structural homology model and identify a previously unappreciated binding pocket present only in GTP bound CDC42 and RHOJ. We use structure-based virtual screening and molecular dynamics simulations^23^ to identify a novel class of compounds that block RHOJ and CDC42 effector interactions, characterize their function *in vitro*, and present proof of principle data that these compounds inhibit tumor growth *in vivo*.

## RESULTS

### Identification of a Drug Binding Pocket Present in GTP bound RHOJ and CDC42

To identify putative drug binding pockets in CDC42 family GTPases, we first examined the crystal structure of CDC42 in both its GDP and GTP bound states. When comparing the crystal structures of GDP-bound CDC42 (PDBid: 1DOA)^24^ and GTP-bound CDC42 interacting with the CRIB domain of PAK6 (PDBid: 2ODB), we discovered a previously unappreciated allosteric pocket at the CDC42-PAK6 protein-protein binding interface that is located on the CDC42 surface in proximity to Ser71, Arg68 and Tyr64 (**Fig. 1A,B**). Specific residues of the structural motifs ‘switch I ’(Val36, Phe37) and ‘switch II ’(Ala59, Tyr64, Leu67, Leu70, Ser71) on the CDC42 surface, define a pocket ∼17 Å away from the GTP binding site, which accommodates the binding of Trp40 on PAK6 (**Fig. 1D**). This pocket is conserved across structures of CDC42 in complex with effectors such as PAK4 (PDBid: 5UPK)^25^, PAK1 (PDBid:1EOA)^26^, and IQGAP (PDBid:5CJP)^27^ -(complete list and structural analyses in **Table S1** and **Fig. S1A,B**). This pocket is also found to be stable during 500ns-long molecular dynamics (MD) simulations of GTP-bound CDC42 (RMSD_pocket_= 2.06 ±0.33 Å, **Fig. S1C**) and is the largest cavity at this protein-protein interface (**Fig. S1E,F**).^28^ This distal pocket was unfolded in the CDC42-GDP complex (**Fig. 1B**).

**Figure 1:**
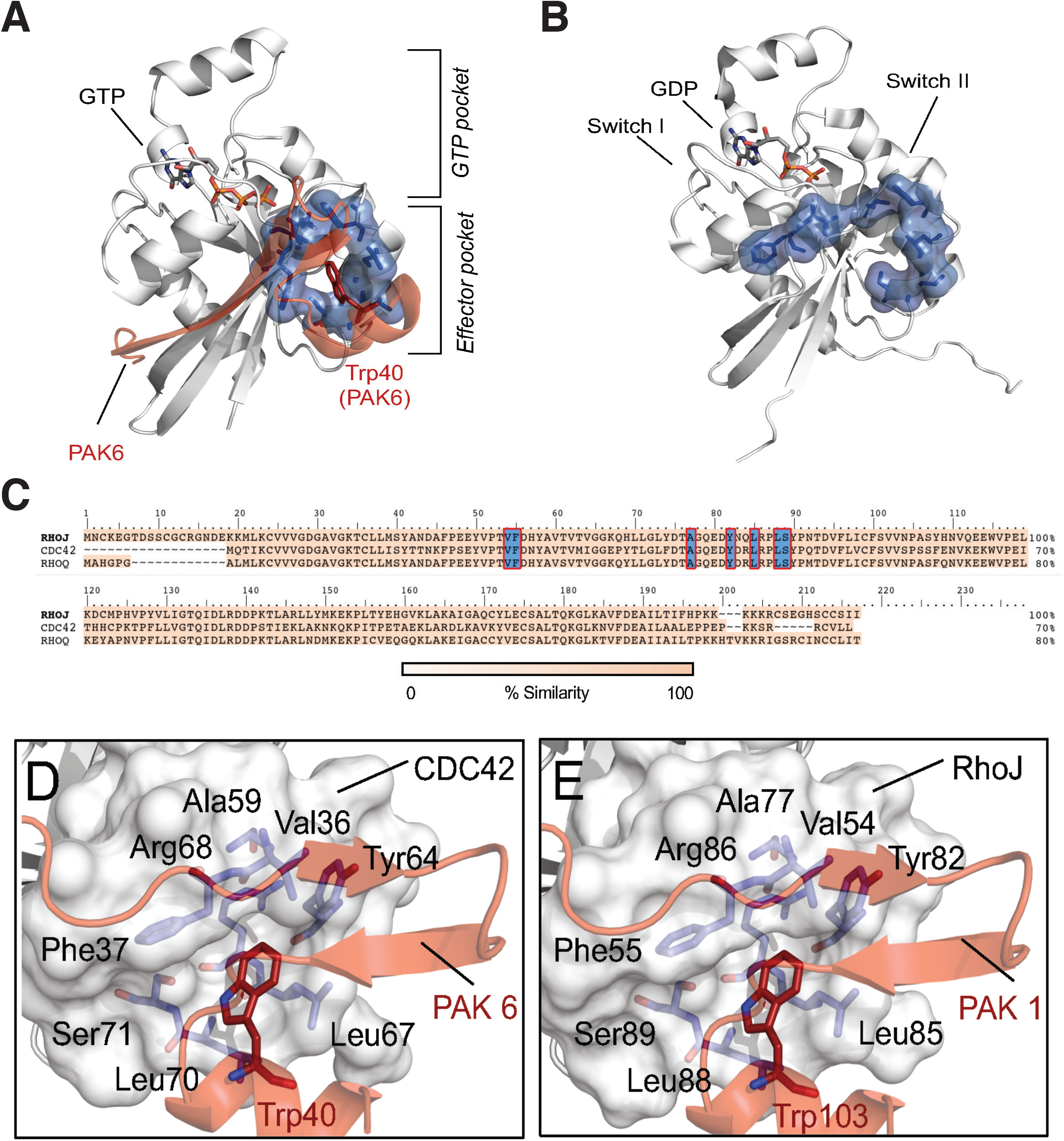
Characterization of a RHOJ/CDC42 allosteric drug binding pocket located at the effector interaction interface. **A)** GTP-bound state of the effector pocket (blue) interacting with Trp40 (red) of PAK6 - PDBid 2ODB. **B)** Unfolded conformation of the effector pocket in the GDP-bound state - PDBid: 1DOA. **C)** Amino acid sequences alignment of RHOJ, CDC42 and RHOQ. The residues defining the drug binding pocket are highlighted in blue. **D)** Close-up view of the CDC42-PAK6 interaction interface. Interactions between Trp40 (red) of PAK6 (transparent red) and the effector binding pocket (light blue) of CDC42 (grey) are shown. **E)** Close-up view of the RHOJ-PAK1 interaction interface. For comparison, modeling of the interaction between Trp103 (red) of PAK1 (transparent red) and the effector binding pocket (light blue) of RHOJ (grey) is shown.

Next, we sought to determine whether this same allosteric pocket could be used to target RHOJ effector interactions. As no crystal structure is available for RHOJ, we built a homology model for RHOJ to use as receptor for a virtual drug screening campaign. The alignment of RHOJ, CDC42, and RHOQ sequences revealed local sequence homology in the domains of RHOJ/RHOQ that interact with its downstream effector PAK, suggesting structural conservation of the pocket (**Fig. 1C-E**). We verified that a similar allosteric binding pocket (involving the interaction of Trp103 of PAK1 with Ser89, Arg86 and Tyr82 of RHOJ)^29^ was structurally conserved (**Fig. 1E**) in RHOJ, and stably maintained through additional 500ns-long MD simulations of GTP-bound RHOJ. We observed that the overall structural fold of RHOJ was preserved during these simulations (RMSD_RHOJ_= 1.20 ±0.14 Å; **Fig. S1D**), as it was the allosteric pocket (RMSD_pocket_= 2.07 ±0.27 Å; **Fig. S1D)**. Thus, this allosteric pocket could be targeted to block RHOJ/CDC42 effector interactions, and is structurally distinct from the CDC42 GEF interaction interface and the K-RAS domains targeted by others (**Fig. S2D-E)**.^30^

### Identification of Putative CDC42 Family Effector Interaction Inhibitors

Next we performed a virtual screening campaign of an internal chemical collection of compounds, which contains a diverse and non-redundant set of ∼20,000 molecules, to identify RHOJ/CDC42 effector inhibitors. Initially, we selected 54 promising compounds and experimentally tested their IC50s in SKMel28 melanoma cells - a cell line that we previously determined was sensitive to RHOJ depletion (**Fig. 2A**).^29^ We then selected compounds with IC50 less than 50 µM (**Table S2**) for further analyses (those with IC50 greater than 50 µM are listed in **Table S3**). We first determined the compounds’ ability to inhibit the interaction between RHOJ or CDC42 and its downstream effector PAK using an established CDC42 interaction assay, which measures the interaction between RHOJ/CDC42 and the PAK-p21 binding domain (**Fig. S2B**). We also measured the kinetic solubility of the active compounds to evaluate their potential for further development. Our initial screening campaign identified ARN12405 as a promising hit (IC_50_ of 16.4 µM in SKM28 cell line and high kinetic solubility of 222 µM) that inhibits RHOJ/CDC42-PAK interactions (**Fig. S2B**). Notably, ARN12405 features a functionalized pyrimidine scaffold bearing a 3-piperidine, a 4-chloro aniline, and a 4-pyridine, respectively, in the 2, 4 and 6 position (**Fig. 2B**). Modeling predicts that ARN12405 fits within the effector pocket of RHOJ and CDC42 (**Fig. S2C**). To further assess the predicted binding poses from our docking results, we performed molecular dynamics (MD) simulations of both RHOJ and CDC42 in complex with ARN12405. As shown in **Figure S2F-G, *Left panels***, our hit compound steadily binds the target pocket throughout the simulations (RMSD_ARN12405_ = 2.00 ± 0.40 Å and 3.24 ± 0.85 Å, for RHOJ- and CDC42-ligand complexes, respectively).

**Figure 2:**
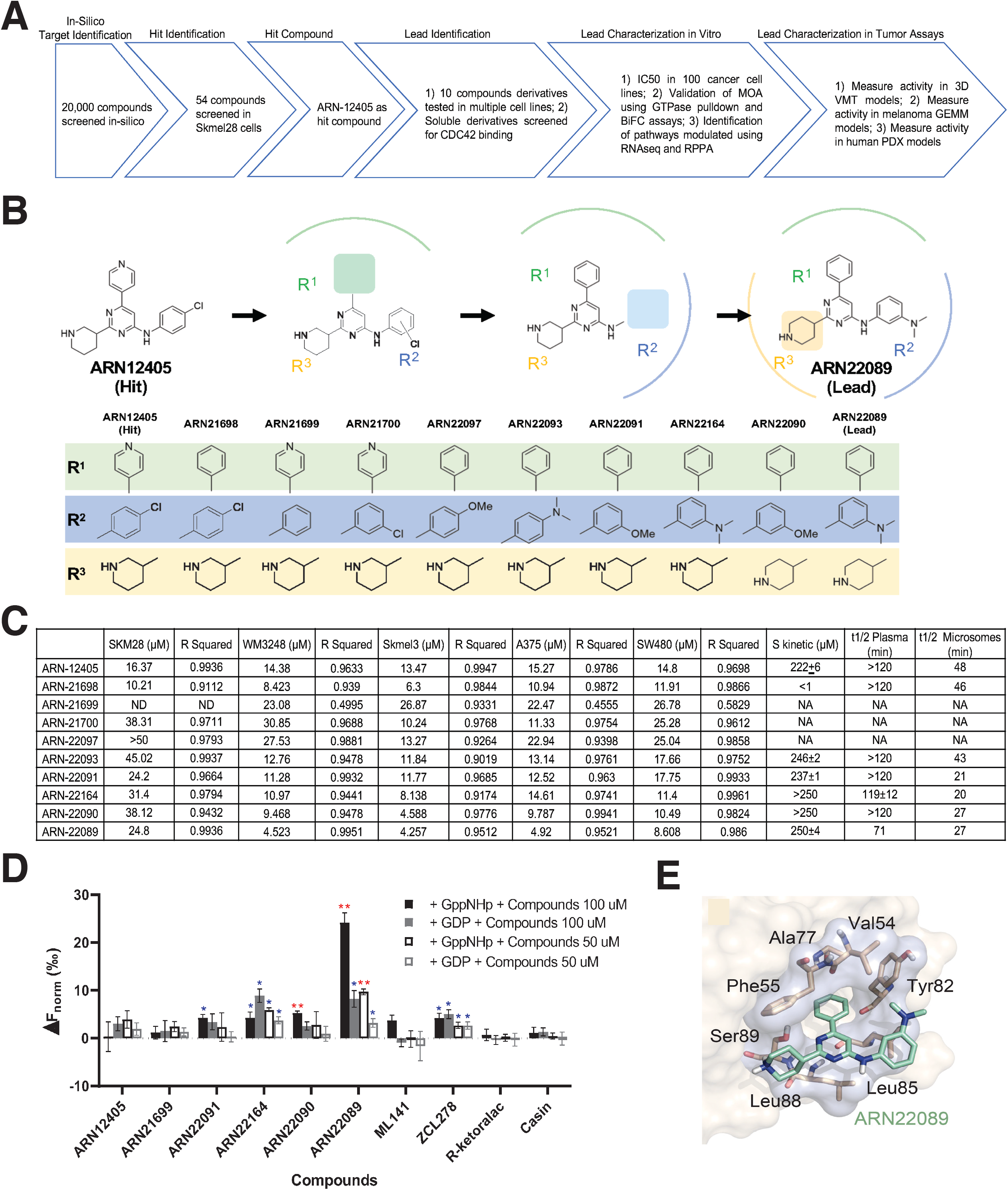
Structure-based screening identifies RHOJ/CDC42 allosteric inhibitors. **A)** Flow diagram of the screening steps to identify RHOJ/CDC42 allosteric inhibitors. **B)** Chemical structures of the hit and synthesized derivatives. Names of the compounds and structures of the indicated R side chains are denoted below. **C)** Table of IC50s, solubilities, and plasma and mouse microsomal half-life of lead and synthesized derivatives. Detailed IC50 curves are shown in **Figure S2A**. **D)** MST was used to screen for binding of the hit, derivatives, and known CDC42 inhibitors to a His-tagged CDC42 fragment. The graph displays the difference in normalized fluorescence (ΔFnorm [%] = ΔFhot/Fcold) at 1.5-2.5 sec between protein:compound sample and a protein only sample at the indicated drug concentrations after loading with the indicated nucleotides. A single asterisk highlights binding with signal to noise ratio equal or higher than 5. Double asterisks are used for binding with signal to noise ratio higher than 12. **E)** Model structure of lead compound bound to the allosteric drug binding pocket of CDC42/RHOJ.

Next we performed hit to lead optimization. We first generated new structural analogues and determined their IC_50_ in five different cancer cell lines (WM3248, Skmel3, A375, SW480, SKM28), as well as kinetic solubility and half-life stability in mouse plasma and in mouse liver microsomes (**Fig. 2A-C,S2A,** synthesis details in Supplemental Information). First, we simplified the starting scaffold to identify the more effective chemical features for RHOJ/CDC42 inhibition, replacing the pyridine heterocycle with a phenyl ring (ARN21698), which slightly increased the activity (IC_50_ 6.3-10.94 μM). Removal or moving the chlorine from para to meta position of the aniline moiety in position 4 (ARN21699, ARN21700), maintaining 4-pyridine, decreased 2 fold the activity compared to the hit in SKM28 cells. Next, we explored additional new aniline substituents while keeping the 3-piperidine heterocycle and the phenyl ring. The replacement of chlorine in para position in ARN21698 by a methoxy (ARN22097) or a dimethyl amino group (ARN22093) resulted in a total or partial loss of activity. In contrast, introducing the same substituents (methoxy and dimethylamino groups) in meta position (ARN22091 and ARN22164) moderately diminished the activity of the compounds. At this point, we moved the piperidine nitrogen from position 3 to 4, keeping either a methoxy or a dimethyl amino substituent in meta position (ARN22090 and ARN22089, respectively – which do not present a chiral center). These compounds are moderately active inhibitors in SKM28 cells (ARN22090 = IC_50_ 38.1 µM, ARN22089 = IC_50_ 24.8 µM). Notably, ARN22089 has a single digit micromolar IC50 activity against the more sensitive cell lines that were tested (WM3248, SKMel3, A375, SW480), an optimal kinetic and thermodynamic solubility (>250 µM and 268 µM, respectively), and a good half-life in mouse plasma (71 minutes) (**Fig. 2C,S5G**). On the considerations of chemical tractability (synthesis and chirality), inhibitory activity, solubility, and half-life, ARN22089 was elected our lead compound. Modeling predicts that ARN22089 fits within the effector pocket of RHOJ and CDC42 (**Fig. 2E,S2D**). For both RHOJ- and CDC42-ARN22089 complexes, the binding pose is preserved during 500ns-long MD simulations (RMSD_ARN22089_ = 3.55 ± 0.62 Å and 2.80 ± 0.61 Å, for RHOJ- and CDC42-ligand complexes, respectively, **Fig. S2F-G**, ***Right panels***). As a further proof of principle, we examined the ability of the hit compound (ARN12405), of other compounds in the chemical class defined above (ARN21699, ARN22090, ARN22091, ARN22164) and of the lead compound ARN22089 to bind to a purified CDC42 fragment. Initial studies by native mass spectrometry determined that purified CDC42 could be effectively loaded with GDP (90% efficiency of loading) or with the GTP analog, GppNHp (>98% efficiency of loading – **Fig. SH-J**). Then, we observed by microscale thermophoresis that our lead ARN22089 binds better to the purified CDC42 as compared to all other compounds in the class (**Fig. 2D**, MST traces in Supplemental Information **1.8-1.9**), and also when compared to other known CDC42 inhibitors (ZCL278, ML141, R-ketoralac and Casin).^31^ Moreover, we found that ARN22089 binds CDC42 preferentially when the protein is the GppNHp loaded state, confirming our modelling results.

### ARN22089 is a Selective CDC42 Family Effector Interaction Inhibitor with Anti-Cancer Activity

To gain a better appreciation of the anti-cancer activity of ARN22089, we started by determining the IC_50_ of this compound in a panel of 100 cancer cell lines. Fifty-five of 100 cell lines that were tested had an IC_50_ less than 10 μM (**Fig. 3A**, **Table S4**), demonstrating a broad spectrum of activity against cancer cells derived from many different tissues. Next, we tested whether our compound was selective for blocking CDC42 family effector interactions in cells. To do this, we tested whether ARN22089 could inhibit the interactions between RHOJ or CDC42 and its downstream effector PAK without affecting the interaction between RAC1 and PAK using an established CDC42 effector assay,^32^ which measures the binding between GTPases and their downstream effectors. In parallel, we performed a similar set of experiments to examine the ability of ARN22089 to inhibit the interaction between less closely related members of the RAS family and their downstream effectors.^33^ WM3248 cells were treated with the indicated doses of ARN22089, cell lysates were prepared, incubated with EDTA to strip GTP and GDP, followed by incubation with GTP or GDP to load the GTPase, and then incubated with PAK1-p21 binding domain (PAK1-PBD) coupled beads or RAF1-RAS binding domain (RAF1-RBD) coupled beads, which were then precipitated and interacting proteins were identified by immunoblotting. ARN22089 inhibited the interaction between RHOJ or CDC42 and PAK1-PBD at 10 μM and 50 μM concentrations only when cell lysates were incubated with GTP (**Fig. 3B**). No interactions were detected between RHOJ or CDC42 and PAK1-PBD when cell lysates were loaded with GDP (**Fig. 3B**), consistent with previously published work.^2^ Notably, ARN22089 did not inhibit the interaction between the most closely related GTPase RAC1 and PAK1-PBD beads or between RAS or RAL and RAF1-RBD beads at either the 10 μM or 50 μM concentrations (**Fig. 3B**), indicating that our lead compound was selective for CDC42 family members and does not inhibit RAC1, RAS, or RAL effector interactions. Our hit compound (ARN12405) similarly blocked RHOJ/CDC42 PAK interaction but not RAC1 PAK interaction (**Fig. S2B**). Taken together, these results indicate that ARN22089 selectively inhibits CDC42 family effector interactions without disrupting signaling from closely related GTPases (RAC1) that are known to be responsible for the cardiotoxicity associated with existing inhibitors of this GTPase family.^20^

**Figure 3:**
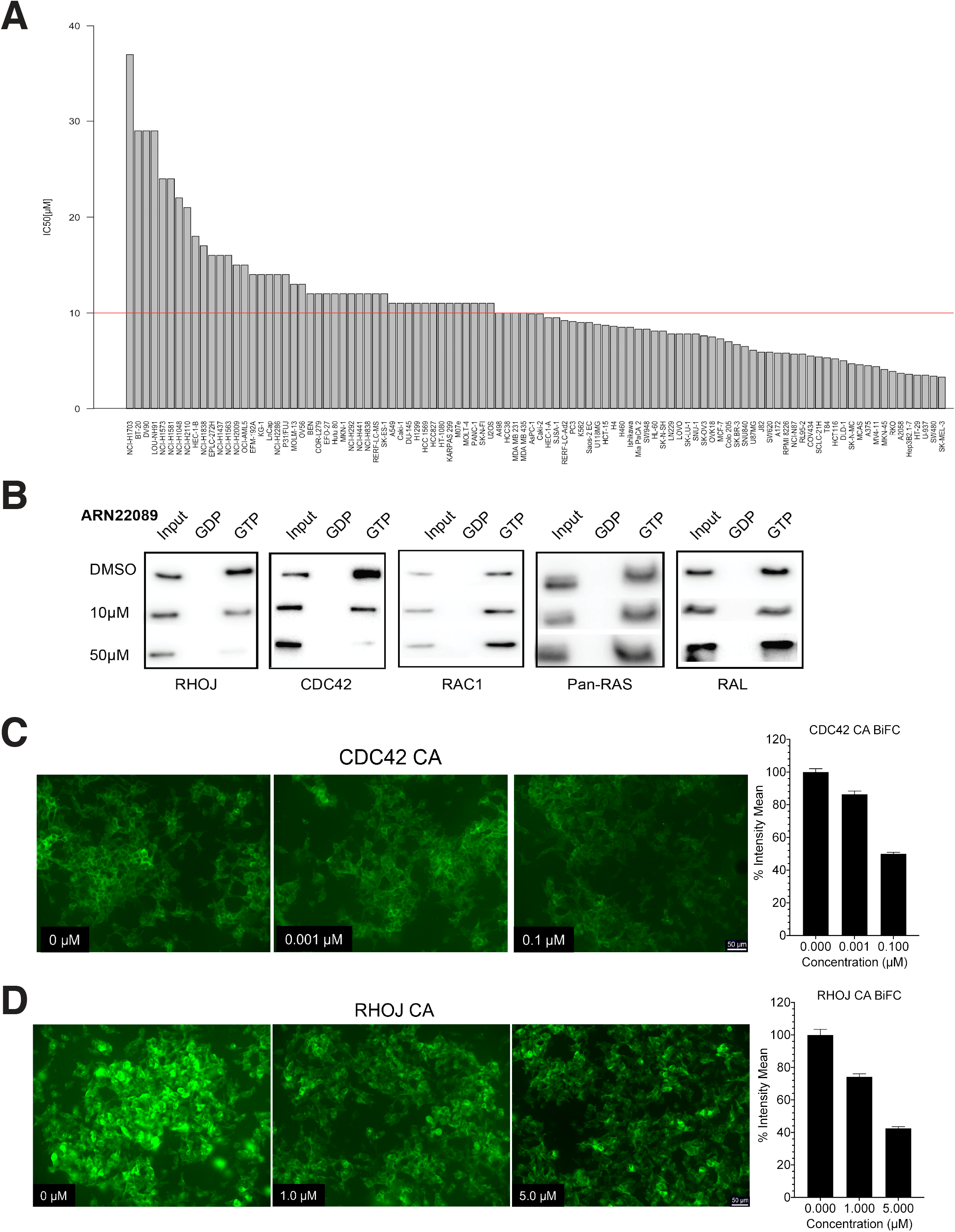
ARN22089 has broad spectrum anti-cancer activity and selectively blocks RHOJ/CDC42 effector interactions. **A)** Bar graph shows IC50 values of ARN22089 in 100 cancer cell lines. **B)** ARN22089 selectively blocks CDC42 effector interactions. WM3248 cells were treated with ARN22089 for 24 hours and lysates were subjected to pulldown assays with PAK1 and RAF1 to measure the ability of the compound to block RHOJ, CDC42, RAC1, RAS, or RAL effector interactions. **(C**) CDC42 and (**D**) RHOJ BiFC cells were treated with 2 µg/mL of DOX to induce CDC42/RHOJ and PAK1 expression and treated with the indicated doses of ARN22089 for 24 and 8 h, respectively. Bar graphs, far right for (**C**) and (**D**), show mean percent GFP intensity for each condition. The 0 µM condition was set to 100%. No DOX condition was used for background subtraction.

To further validate that ARN22089 inhibits CDC42 effector interactions, we examined whether ARN22089 could inhibit CDC42-PAK interactions in cells. To measure CDC42 effector interactions in cells, we used a BiFluorescence complementation (BiFC) assay,^34^ where CDC42 or RHOJ was linked to the N-terminus of Venus fluorescence protein (FP) fragment, while PAK1 is linked to a C terminal Venus FP fragment (**Fig. S3A**). We then measured CDC42/RHOJ PAK interactions by measuring YFP fluorescence in cells. Initial optimization studies revealed that we could visualize GFP indicative of CDC42 PAK interactions in cells that expressed constitutively active (CA) forms of RHOJ/CDC42 and PAK in the presence but not absence of Doxycycline (DOX) which induces the transcription of both proteins (**Fig. S3B**). We next determined the percent intensity of CDC42-PAK interactions using different doses of ARN22089. We observed that ARN22089 could inhibit CDC42-PAK interaction as measured by our BiFC assay with an estimated EC50 of 100 nM (**Fig. 3D, S3C**), consistent with the observation that our hit compound bound to the CDC42 fragment with comparable affinities to compounds with nanomolar affinities (**Fig. 2D**).^1^ We observed that ARN22089 could inhibit RHOJ/PAK interactions at an approximate EC50 between 1-5 μM (**Fig. 3E, S3D**), consistent with the IC50s observed in cancer cell lines tested.

### ARN22089 Inhibits MAPK and S6 Phosphorylation and activates NFkB signaling *in vitro*

Published studies indicate that CDC42 GTPases are known to activate a spectrum of signaling cascades. GTP bound CDC42 family GTPases bind to and activate p70S6K.^35^ Active p70S6K goes on to phosphorylate the S6 ribosomal protein at two different locations, 234/235 and 240/244.^36^ GTP bound CDC42 family members also activate PAK kinases, whose downstream targets include members of the MAPK and NFkB pathways.^37^ To understand the effect of ARN22089 on cells, we first sought to examine how drug treatment influenced the activation of kinase signaling pathways in melanoma cells. WM3248 melanoma cells were treated with 5, 10, or 20 μM ARN22089 for six hours, cell lysates were prepared, and proteins were hybridized with a reverse phase protein array consisting of 486 antibodies to quantitatively examine how the drug influenced protein abundance and protein phosphorylation. This analysis revealed that the compounds were fairly selective, as the abundance of 38 proteins and the phosphorylation of 10 proteins were significantly affected at both the 10 and/or 20 μM doses (**Fig. 4A**, *Left*, **Table. S7**). We observed that ARN22089 could inhibit S6 phosphorylation at both serine 235/236 and 240/244 residues with both 10 and 20 μM doses (**Fig. 4A**, *lower inset*). We also observed that ARN22089 could significantly inhibit phosphorylation at the 235/236 sites in both WM3248 and A375 melanoma cells with 10 and 20 μM doses at 6 hours (**Fig. 4B**) and with a dose of 5 μM at 24 hours (**Fig. 4B**). Another phosphorylation event that was inhibited by ARN22089 was MAPK1 (aka ERK). We observed that ARN22089 inhibited ERK phosphorylation with 10 and 20 μM doses at 6 hours (**Fig. 4B**) and with a 5 μM dose at 24 hours in A375 and WM3248 cells (**Fig. 4B**). Taken together, these studies indicate that ARN22089 is a selective inhibitor of S6 and ERK phosphorylation *in vitro*.

**Figure 4:**
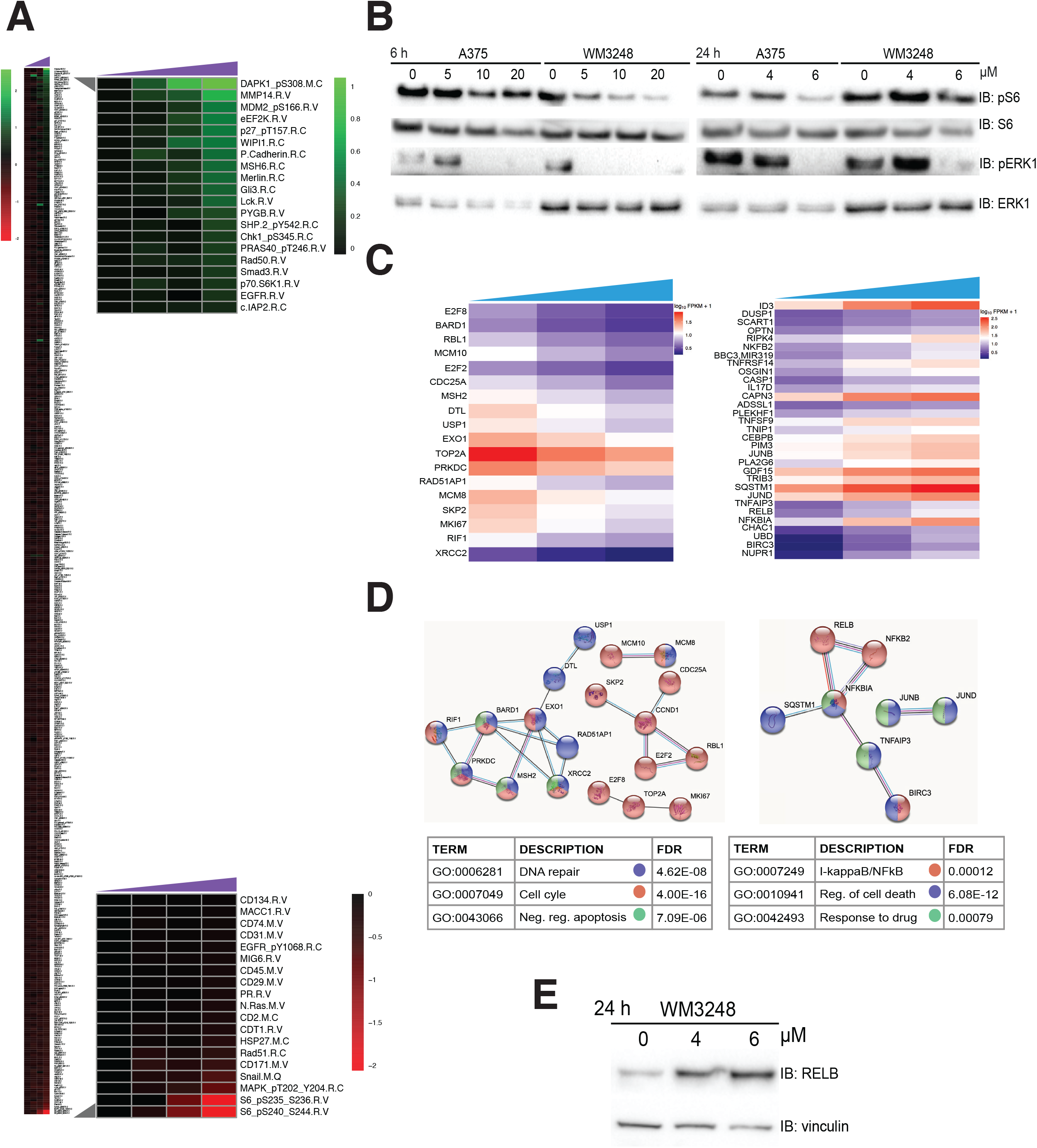
Characterization of the activity of ARN22089 in melanoma cells. **A)** RPPA heatmap showing effects of drug treatment on protein accumulation/phosphorylation. Colors represents values of each treated dose normalized to vehicle (0 µM) (see **Table S6**). Left, heatmap shows results for 486 proteins assayed. Right, insets include proteins that were significantly modulated by drug treatment (see **Table S7**). **B)** WM3248 or A375 cells were treated with the indicated doses of ARN22089 and the accumulation of pS6 and pERK (targets significantly modulated in the RPPA analysis) were measured by immunoblotting. **C)** Profile heatmaps of mRNA sequences that were differentially transcribed after drug treatment (cutoff q-value < 0.05, **Table S8**). **D)** Protein-protein interactions networks modulated by drug treatment. Significant genes shown in (**C**) were subjected to STRING analysis to identify protein-protein interaction networks. Nodes are color coded based on Gene ontology analysis shown in the table below. **E)** Immunoblot for RELB from WM3248 cells that were treated with the indicated doses of ARN22089 for 24 h.

Next, we sought to take a broader view of the pathways engaged by ARN22089. To approach this, we treated cells with either 0, 4, or 6 μM of ARN22089, harvested the cells, and performed RNA sequencing to identify transcripts that were up- or down-regulated upon ARN22089 treatment. We observed that ARN22089 treatment induced the expression of genes involved in cell death and modulated NFκB signaling (**Fig. 4C,D**, **Table S8**), consistent with the known role of PAK kinases in regulating NFκB signaling ^34^. We showed that the RELB protein level is increased with drug treatment compared to untreated cells, corroborating our RNA-seq analysis (**Fig. 4E**). Taken together, these results suggest that ARN22089 inhibits two known CDC42 effector functions-p70S6K activation and PAK activation.

### ARN22089 Specifically Inhibits Tumor Angiogenesis in 3D Vascularized Microtumors

CDC42 and RHOJ are both known to have specific roles in tumor angiogenesis. To better understand how ARN22089 impacts tumor angiogenesis, we sought to test the ability of these compounds to inhibit vessel formation around tumors. Here we utilized a vascularized microtumor platform (VMT) - a “tumor-on-a-chip” platform - that incorporates human melanoma cells, which are grown in a 3D extracellular matrix (ECM), and delivers nutrient to the cells via perfused micro-vessels^38–41^ (**Fig. S4A-C**). Each chamber of the VMT were loaded with mCherry endothelial cells, GFP labeled tumor cells (A375 and WM3248), fibroblasts, and pericytes. After 4 days, these cells formed a capillary bed that could feed the growing tumor, similar to newly formed vessels *in vivo* (**Fig. S4D**). Nutrients and drug (ARN22089, FRAX597, vehicle) were perfused into the chamber from the high pressure “arterial side” and traversed to the tumor through the vascular network every two days at the indicated doses. The effects of each compound on the number of GFP tumor cells and the length of endothelial vessels was measured. Treatment of VMTs at a concentration of 2 μM ARN22089 inhibited the growth of both A375 cells and the mCherry labeled blood vessels around the tumor (**Fig. 5A**). In contrast, 2 μM FRAX597 did not significantly inhibit the growth of the A375 cells or the blood vessels (**Fig. 5A**). These findings were confirmed in WM3248 cells in which a dose of 2 μM ARN22089 caused significant tumor and vessel regression compared to both control and FRAX treated VMT (**Fig. 5B**). These observations indicate that ARN22089 inhibits both tumor growth and angiogenesis.

**Figure 5:**
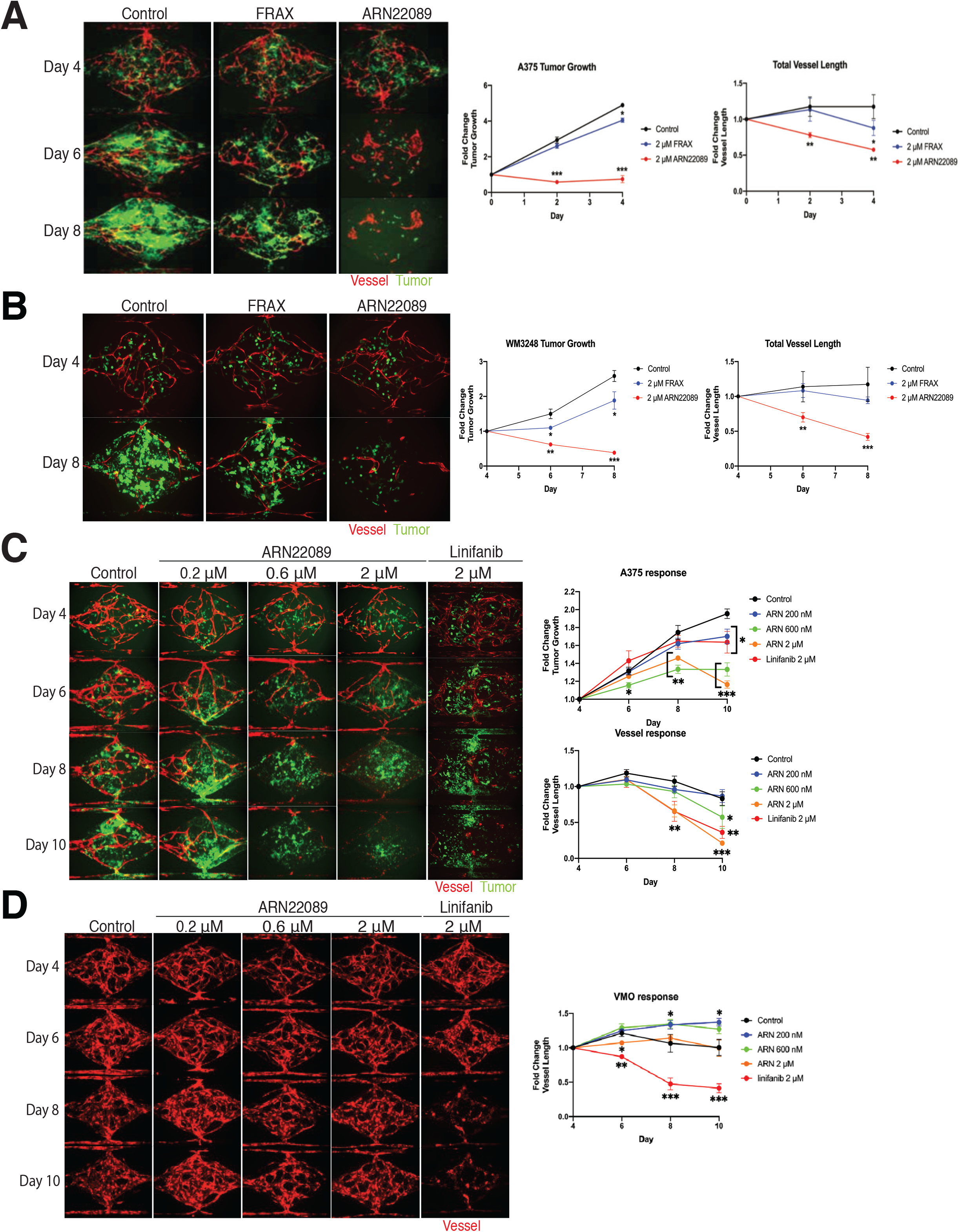
ARN22089 inhibits tumor growth in a Vascularized Microtumor (VMT) model. Representative micrographs of **(A)** A375 or **(B)** WM3248 VMTs (organoids that contain cancer cells and endothelial cells) treated with control, 2 µM FRAX or ARN22089. Quantification of tumor cell growth and vessel length shown on right. **C)** Micrographs VMT of WM3248 treated with increasing concentration of ARN22089, corresponding quantification shown on right. The angiogenesis inhibitor Linifanib at 2 µM is shown as a comparison. **D)** Micrographs of VMOs (organoids with only endothelial cells) treated with increasing concentrations of ARN22089 or Linifanib and corresponding quantification of vessel length is shown.

Next, we sought to more closely examine whether ARN22089 was a specific inhibitor of tumor angiogenesis. We generated VMTs that contained A375 cells and endothelial cells and VMOs (vascularized micro-organs) that contained vessels but no tumor cells. We observed that ARN22089 could inhibit the growth of both tumor cells and vessels in VMTs at a 600 nM and 2 μM doses, significantly more effective than the angiogenesis inhibitor linifanib (**Fig. 5C**). In contrast, ARN22089 did not inhibit the growth of VMOs (**Fig. 5D**). We observed that linifanib, in contrast, did inhibit angiogenesis in VMOs (**Fig. 5D**). These results suggest that ARN22089 specifically inhibits tumor and not normal angiogenesis, consistent with the selective role of the CDC42 family member RHOJ in tumor angiogenesis.^6^

### ARN22089 Has Drug-Like Properties and Inhibits Tumor Growth *in vivo*

Once we determined that our RHOJ/CDC42 interaction inhibitor had a broad spectrum of activity *in vitro*, we next determined whether it has drug-like properties. First, we verified that ARN22089 has no significant off-target effects as agonist or antagonist for a panel of 47 classical pharmacological targets, even at concentrations up to 25 μM (**Fig. S5A-E**). Notably, ARN22089 does not target the hERG channel (**Fig. S5C**), which in other cases has been an impediment in developing safe drugs.^42^ We then determined the pharmacokinetics of ARN22089 in experimental animals after intraperitoneal, intravenous, and oral administration (**Fig. 6A**). The compound was well tolerated in experimental animals and had drug-like PK properties (**Fig. 6A, S5F**). In addition, we incubated the compound with both human and rat liver microsomes and determined that the compound had a half-life of 107 minutes after incubation with rat microsomes and 510 minutes after incubation with human microsomes (**Fig. S5G**).

**Figure 6.**
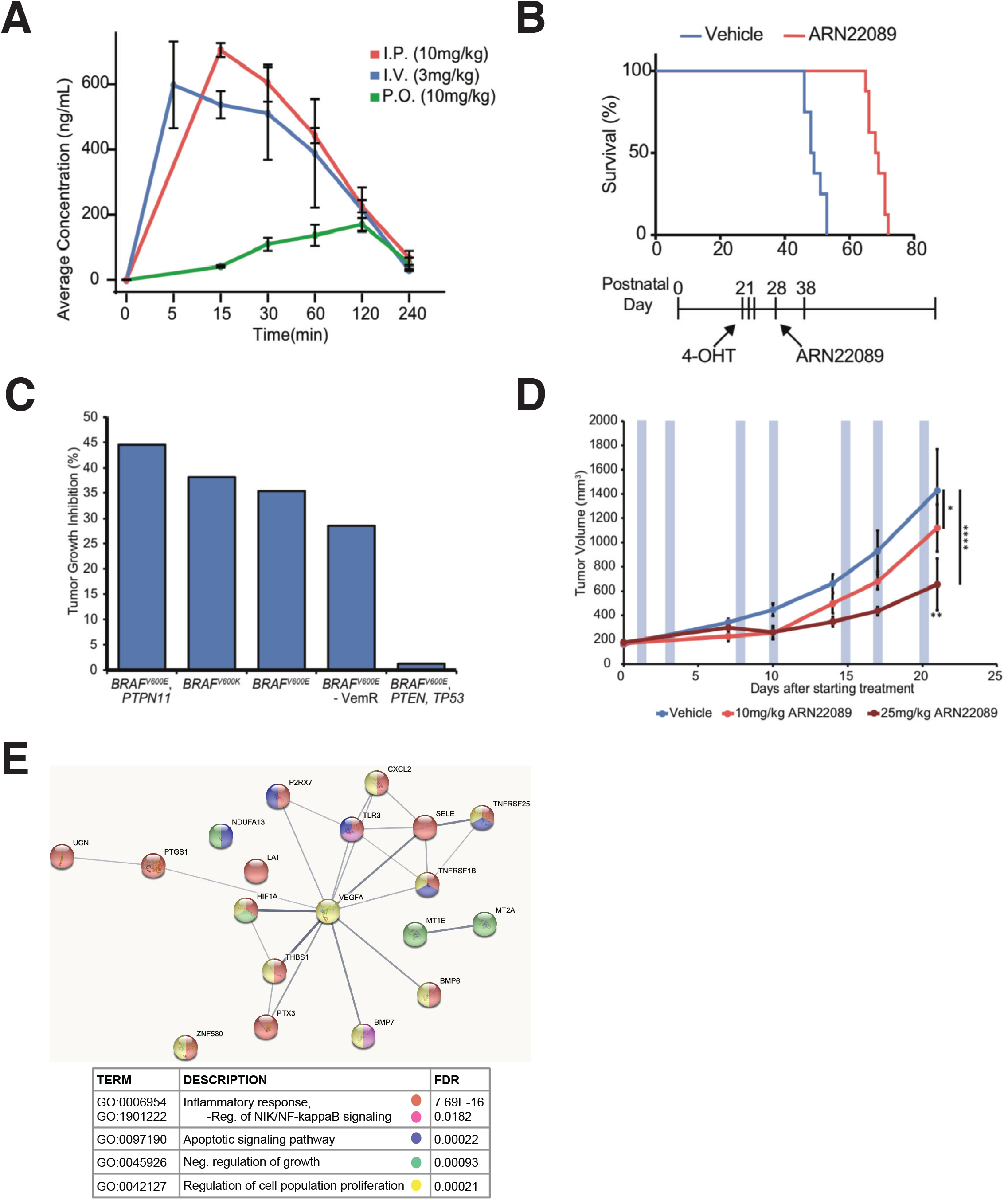
ARN22089 inhibits tumor growth in mouse models. **A)** ARN22089 delivered at 10mg/kg I.P. and 3mg/kg I.V. has drug like properties. PK of compounds was calculated as described in the methods. Complete data is shown in Fig S5F. **B)** ARN22089 inhibits melanoma development in a murine *BRAF^V600E^, PTEN^flox/flox^, Tyr:Cre^ERT2^* model. Melanoma was induced at P21-23 with topical treatment of 25mg/mL 4-OHT. Mice were then treated with vehicle (blue) or 10mg/kg I.P. ARN22089 (red) for 10 days between P28-P38. **C)** ARN22089 inhibits tumor growth in *BRAF* mutant PDX tumor models (listed by verified mutations). Mice were treated for 14 days with 10mg/kg I.P. ARN22089 starting when PDX tumors reached 150-200mm^3^. Tumor growth inhibition was compared with matched vehicle treated mice (see also Figure S6A,B). **D)** ARN22089 inhibits tumor growth in a dose responsive manner. *BRAF^V600E^, PTPN11* PDX tumors were treated with vehicle (blue), 10mg/kg ARN22089 (light red), or 25mg/kg ARN22089 (dark red) twice weekly after tumors reached 150-200mm^3^, blue shaded boxes denote I.V. treatments (see also **Figure S5**). **E)** RNA from the tumors described in (**D**) were sequenced. STRING was used to identify differentially expressed protein-protein interaction networks. Nodes are color coded based on Gene Ontology indicated in the table below.

To test the efficacy of the compound *in vivo*, we first induced BRAF mutant melanoma tumors with topical tamoxifen in *Tyrosinase::CreERT2; Braf^CA/+^; Pten^fl/fl^;RhoJ ^+/+^* animals at P21 ^2^, and treated the mice with ARN22089 for 10 days using a BID, IP dosing regimen. Inhibitor treatment prolonged the survival of tumor carrying mice (mean survival: control 51.5, treated 68.5) (**Fig. 6B**) after only 10 days of treatment. Once we had established that this compound could inhibit the growth of mouse tumors in immunocompetent mice, we next examined the ability of this compound to inhibit the growth of patient-derived xenografts (PDXs) in NOD scid gamma (NSG) mice. Animals were inoculated with PDX tumors, and tumor treatment was initiated after tumors reached a size of 150-200 mm^3^. Animals were treated with 10 mg/kg IP BID for two weeks, after which tumors were allowed to continue to grow until they reached endpoint (1500-2000 mm^3^). The compound significantly inhibited the growth of 2/5 PDX tumors tested as measured by % tumor growth inhibition and generally inhibited the growth of 4/5 of the tumors tested (**Fig. 6C,S6A**). Interestingly, the PDX models tested included one PDX model that was derived from a patient which had failed both a BRAF inhibitor and immunotherapy, also responded to the treatment (**Fig. S6B**). Histologic evaluation of treated tumors revealed that ARN22089 treatment for 2 weeks increased the amount of tumor necrosis in three of the four tumors that responded to drug (**Fig. S6C,D**). To gauge the effects of long-term ARN22089 treatment on tumor growth, we inoculated mice with one BRAF mutant PDX that was known to have consistent growth properties, and began treating the mice with ARN22089 with 0, 10 mg/kg, or 25 mg/kg IV BIW until control tumors reached endpoint. ARN22089 inhibited tumor growth in a dose responsive manner, without modulating weight of mice (**Fig. 6D,S6E**). We harvested tumors treated at the 0 and 25 mg/kg IV dose and harvested RNA from tumors. RNA-seq analysis of tumors revealed that ARN22089 treatment modulated NFκB signaling in 25 mg/kg IV treated tumors, as was observed in *in vitro* treated cells (**Fig. 6E**, **Table S9**). Taken together, these results provide proof of principle that ARN22089 can inhibit tumor growth and vasculogenesis *in vivo* by modulating similar pathways that were observed *in vitro*.

## DISCUSSION

Targeting CDC42 family GTPases has been difficult secondary to its globular structure and limited small molecule binding pockets,^1^ its extensive post-translationally modification in cells,^43^ and the fact that these proteins are localized to discrete nanodomains^44^. We took a multipronged approach to identify molecules that would inhibit the protein-protein interaction between CDC42 and its downstream effectors. We identify a structurally conserved, allosteric drug binding pocket that is folded in GTP bound CDC42 and RHOJ, is stable through 500 ns-long molecular dynamic simulations, and is only partially formed and solvent exposed in GDP bound CDC42 (**Fig. 1**). This pocket directly interacts with Trp40 of PAK, indicating that molecules targeting this pocket would inhibit RHOJ/CDC42 and PAK interactions.

Starting from the discovery of the initial hit compound ARN12405, we built a structure-activity relationship (SAR) study that allowed us to develop the most promising new molecular entity, ARN22089, which we further characterized *in vitro*. First, we verified that: 1) the lead (ARN22089) could bind to CDC42; 2) it bound preferentially to active CDC42; 3) it bound more avidly than other known CDC42 inhibitors (**Fig. 2D**). Second, we examined whether ARN22089 has selective activity against CDC42 GTPases. ARN22089 could inhibit the interaction between RHOJ/CDC42 and PAK but did not block the interactions between the closely related RAC1 GTPase and PAK or RAS/RAL and Raf at doses as high as 50 μM (**Fig. 3B**). As a further validation, we established a bifluorescence complementation assay, which has been used to measure RHOJ protein-protein interactions,^34^ to measure whether our compound could inhibit the protein-protein interaction between RHOJ/CDC42 and PAK1. ARN22089 could inhibit the interaction between RHOJ/CDC42 and PAK1 with an EC50 in the nanomolar range for CDC42 and the uni-digit micromolar range for RHOJ (**Fig. 3C,D**).

Many groups have sought to develop small molecules that inhibit protein-protein interactions,^45^ including between Ras family members and their downstream effectors.^46,47^ While several of these agents have made it to the clinic, others failed secondary to their suboptimal selectivity and off target-effects.^4^ ARN22089 had single digit micromolar EC50s in a panel of cell lines (**Fig. 3A**) consistent with the EC50 values observed with other RAC/CDC42 family inhibitors^48^ and non-covalent RAS family protein-protein interaction inhibitors.^46,47^ Using a reverse phase protein array, we observed that ARN22089 selectively modulated signaling pathways known to be downstream of CDC42 with little off target effects on other kinase cascades-treatment of cells with 10 or 20 μM ARN22089 significantly modulated the expression of only 38 targets and the phosphorylation of only 10 targets (**Table S7**). CDC42 can directly bind and activate p70S6K,^35^ inducing the phosphorylation of S6K at 234/235. This specific phosphorylation event was inhibited by ARN22089 (**Fig. 4A**). Moreover, we saw a more potent effect on this phosphorylation event when melanoma cell lines were incubated with this compound for longer times (**Fig. 4B**). PAK kinases can modulate ERK signaling by phosphorylating MEK at Ser298^49^ and also by phosphorylating Raf-1.^50^ We observed that ARN22089 could inhibit ERK phosphorylation (**Fig. 4A,B**) and block the phosphorylation of MEK at Ser298 (data not shown). Taken together, these data indicate that while ARN22089 has a similar affinity for the protein-protein interaction interface as other drugs, it is highly selective as we observed: 1) no off-target activity as agonist or antagonist against a panel of 47 classical pharmacological targets (**Fig. S5A-E**); 2) a high degree of on-target modulation of CDC42 signaling (**Fig. 4**).

Several other lines of evidence support the development of ARN22089 as a treatment for cancer. ARN22089 shows excellent solubility and has good chemical and metabolic stability, has a favorable absorption-distribution-metabolism-excretion (ADME) profile *in vitro,* and has favorable pharmacokinetics *in vivo* (intraperitoneral, intravenous, and oral administration). CDC42 family members regulate not only tumor cell proliferation but also tumor angiogenesis.^5,6,14,40^ ARN22089 inhibited not only tumor cell growth but also inhibited vessel elongation in three dimensional vascularized tumor models, in contrast to PAK inhibitors, which affect tumor proliferation but not vessel growth (**Fig. 5A-C**). Anti-angiogenic effects were specific for tumor-associated vessels, as ARN22089 had no impact on vessel growth when tumors were not present (**Fig. 5D**). Finally, ARN22089 modulated the expression of genes involved in inflammation and apoptosis (**Fig. 4D,6E**), consistent with a role for the drug in inducing inflammation and apoptosis in tumors.^51^

Before detailed formulation development, toxicokinetic, and pharmacokinetic characterization of the compound, we wanted to get a sense of the *in vivo* anti-cancer activity of the compound using a crude formulation. We demonstrated that short term treatment of mice with ARN22089 could prolong survival in a BRAF mutant autochthonous mouse model of melanoma (**Fig. 6B**). In addition, short term treatment could slow the growth of 4/5 BRAF mutant human PDXs to some extent (**Fig. 6C**), including one PDX model that was generated from a BRAF inhibitor resistant tumor (**Fig. S6B**). Drug treatment induced necrosis in treated tumors, a phenomenon that is often observed with angiogenesis inhibitors ^52^, consistent with the data obtained in the VMT models (**Fig. S6C,D**). Notably, prolonged treatment of tumors with increasing doses of ARN22089 induced more profound tumor growth inhibition, an effect that was dose responsive (**Fig. 6D**). Although further optimization of dosing is still needed, gene expression analysis of tumors revealed that ARN22089 modulates many of the same pathways *in vivo* that were observed *in vitro*. Given the unique spectrum of pathways modulated by ARN22089 and the effects of the drug on the tumor and its microenvironment, this therapy has logical applications both as a primary treatment, as a combination treatment to reduce the dose limiting toxicities of existing agents, or as a treatment for tumors that have developed resistance to other agents. In a broader context, these studies provide a roadmap for the rational structure-based design of drugs targeting other RHO superfamily GTPases for other applications.

## MATERIALS AND METHODS

### Computations

#### Homology Modeling

As a template structure, we employed the CDC42 protein in complex with the CRIB domain of Pak6 (PDB code 2ODB, resolution of 2.4 Å). By means of Prime (Jacobson et al 2004) software implemented in Maestro, we modeled the FASTA sequence of RHOJ on the 2ODB X-ray structure (i.e., Cdc42 template), and then the resulting structure was refined by using the Protein Preparation Wizard (Sastry et al 2013) workflow implemented in Maestro. According to this procedure, hydrogen atoms were added, and charges and protonation states were assigned titrating the protein at physiologic pH. The steric clashes were relieved by performing a small number of minimization steps, until the RMSD of the non-hydrogen atoms reached 0.30 A.

#### Virtual screening

In the CDC42-PAK6 complex, we identified a pocket on the CDC42 surface in proximity of Ser71, Arg68 and Tyr64. Since this surface cavity is conserved in RHOJ, displaying the same residues as in CDC42 (i.e. Ser89, Arg86 and Tyr82), and also the tryptophan is conserved in PAK1, we used this pocket to center the grid. The cubic grid box was centered on Ser89 of RHOJ, having a dimension of 26 × 26 × 26 Å3. We screened a set of nonredundant ∼20,000 molecules. These molecules belong to a proprietary chemical collection available in the D3 Department at IIT. The molecules ‘database was prepared using LigPrep software implemented in Maestro. Firstly, we added hydrogens and generated ionization states at pH 7.4 ± 0.5. Then, we generated tautomers and all stereochemical isomers. For each structure containing a ring moiety, the low-energy conformation was computed and retained. Lastly, a short minimization step was carried out to relax the 3D structure of each molecule. At this point, we filtered the resulting database to discard molecules that are not endowed with drug-like properties. To do so, we firstly computed the ADME descriptors for each molecule using QikProp software implemented in Maestro. As filter, we discarded all the molecules that do not respect the Lipinsky’s rule of five.^53^ We used Glide to perform the virtual screening, using Single Precision and retaining one pose for each ligand.

#### Molecular Dynamics Simulations

Molecular dynamics (MD) simulations were performed considering both our RhoJ structural model and Cdc42 X-Ray structure (PDBid 2ODB) in their ligand-free state, as well as four different protein-ligand complexes as obtained from our docking calculations. Specifically, two systems were built using our RhoJ structural model bound to either ARN22089 or ARN12405. Other two systems included Cdc42 (PDBid 2ODB) in complex with either ARN22089 or ARN12405. Additionally, the GTP substrate as well as the catalytic Mg^2+^ ion are present at the active site of the proteins, in all the systems. These models were hydrated with a 14Å layer of TIP3P water molecules^54^ from the protein center. The coordinates of the water molecules at the catalytic center were taken from PDBid 2ODB. Sodium ions were added to neutralize the charge of the systems. The final models are enclosed in a box of ∼89·89·89 Å3, containing ∼18,500 water molecules, resulting in ∼ 59,000 atoms for each system.

The AMBER-ff14SB force field^55^ was used for the parametrization of the protein. The parameters for the ligands ARN12405 and ARN22089 were determined via Hartree-Fock calculation, with 6-31G* basis set, convergence criterium SCF=Tight after structure optimization (DFT B3LYP functional; 6-31G* basis set). Merz-Singh-Kollman scheme^56^ was used for the atomic charge assignment. The GTP and the Mg^2+^ were parametrized according to Meagher KL et al.^57^ and Allner et al.^58^ respectively. Joung-Chetham parameters^59^ were used for monovalent ions.

All MD simulations were performed with Amber 20^60^ and all the systems were object of the following equilibration protocol. To relax the water molecule and the ions, we performed an energy minimization imposing a harmonic potential of 300 kcal/mol Å2 on the backbone, the GTP and the docked compound, when present. Then, two consecutive MD simulations in NVT and NPT ensembles (1 ns and 10 ns, respectively) were carried out, imposing the previous positional restraints. To relax the solute, two additional energy minimizations steps were performed imposing positional restraints of 20 kcal/mol Å2 and without any restraints, respectively. Such minimized systems were heated up to 303 K with four consecutive MD simulations in NVT (∼0.1 ns, 100 K) and NPT ensembles (∼0.1 ns, 100 K; ∼0.1 ns, 200 K; ∼0.2 ns, 303 K), imposing the previous positional restraints of 20 kcal/mol Å2. We used the Andersen-like temperature-coupling scheme^61^ while pressure control was achieved with Monte Carlo barostat at reference pressure of 1 atm. Long-range electrostatics were treated with particle mesh Ewald method. We performed an additional MD simulation (∼1.5 ns) in the NPT ensemble at 303 K without any restraint to relax the system at such temperature. Finally, multiple replicas of 500 ns were performed in the NPT ensemble for each system with an integration time step of 2 fs.

### His-Cdc42 production and purification

His-Cdc42 wild-type (aminoacids Ile4-Pro182) was expressed in *E. coli* BL21 (DE3) cells using the pET28a expression vector. Overexpression was induced by 0.1 mM IPTG at OD_600_ 0.8 in Luria Bertani broth and incubated over night at 18 °C. Cells were harvested by centrifugation at 6000 xg for 30 min and the pellet was stored at – 80 °C. Cells were defrosted by incubation at room temperature in 50 mM Tris-Cl buffer pH 7.5, 400 mM NaCl, 5 mM MgCl_2_, 50 μM GDP, 40 mM imidazole, 10 μg/ml DNAse (Sigma) and 30 µg/ml Lysozyme (Sigma). Cells were lysed by sonication and centrifuged at 43000 xg at 4 °C for 1 h. The supernatant was incubated for 3h at 4 °C, rotating, with Ni-NTA resin prior in batch purification. His-Cdc42 was eluted with 300 mM imidazole.

### Preparation of GppNHP/GDP-bound GTPase

For the loading of GDP, the purified protein was dialyzed over night at 4 °C in 50 mM Tris pH 8.5, 200 mM ammonium sulfate, 50 µM GDP (Jena Bioscience). For the loading of GppNHp, His-Cdc42 was instead dialyzed with 20 µM GppNHp (Jena Bioscience) in the presence of 5 U of Quick-CIP alkaline phosphatase (New England Biolabs). After dialysis, 2 mM MgCl_2_ was added to the solution to stabilize nucleotide binding. The two samples were buffer exchanged in 20 mM Hepes pH 7.5, 40 mM NaCl, 5 mM MgCl_2_, 1 mM DTT and loaded on a RESOURCE Q (Cytiva) column for anion exchange chromatography. The efficiency of nucleotide loading was evaluated by native state mass spectrometry.^62^ The purified proteins were buffer exchanged in 10 mM ammonium acetate pH 6.8 and diluted to 3 μM in 10 mM ammonium bicarbonate pH 6.5 added with final 3% acetonitrile. The samples were then infused at 40 µL/min in an electrospray ion source, coupled to a Synapt G2 QToF mass spectrometer operating in positive ion mode. Spectra were acquired over the 500-4000 m/z range.

### Target binding by microscale thermophoresis

MicroScale Thermophoresis (MST) experiments were performed according to the NanoTemper technologies protocols in a Monolith NT.115 Pico (Pico Red / Nano Blue - NanoTemper Technologies). His-Cdc42 affinity for the RED-tris-NTA label was not optimal therefore Alexa Fluor 647– NHS dye was used. His-Cdc42 was labeled following the instructions of the MO-L001 Monolith NT Protein Labeling Kit RED – NHS (NanoTemper Technologies). Labelled protein concentration in the binding reactions was 10 nM while compounds concentration was either 50 or 100 μM. DMSO concentration was maintained constant across samples at either 0.5 % or 1 %. Solutions were prepared in 100 mM Trizma® base (Sigma) pH 7.5, 40 mM NaCl, 0.05% v/v Tween 20 and incubated 5 min before loading on Premium Capillaries and analysis. Binding was detected at 24 °C, MST power high and 20% LED power. The MST traces were recorded as follows: 3 s MST power off, 20 s MST power on and 1 s MST power off. The difference in normalised fluorescence (ΔF_norm_ [%]= F_hot_/F_cold_) between protein:compound sample and a protein only sample at 1.5-2-5 sec is calculated and plotted through MO.Affinity analysis v2.3 (NanoTemper Technologies) and GraphPad Prism 8.0.0 (GraphPad Software, San Diego, California USA). Signal to noise ratio was used to evaluate the quality of the binding data according to NanoPedia instructions (NanoTemper Technologies). Only a signal-to-noise ratio of more than 5 was considered acceptable while a signal to noise of more than 12 was considered excellent.

### CDC42 interaction assay

A CDC42 activation assay was performed according the manufacturer’s protocol (Cell Biolabs, San Diego, CA) as described previously ^2^. Briefly, cells expressing high RHOJ (WM3248 or WM983B) were treated with ARN22089 at the indicated doses. 24 hours later the lysates from treated and untreated cells were unloaded of guanosine nucleotides and either loaded with GDP or GTPγS. Agarose beads conjugated with the PAK1 PBD or RAF RBD were used in pull down assays for RHOJ, CDC42, RAC1 or RAS and RAL, respectively. Precipitated lysates were then immunoblotted with indicated antibodies.

### Immunoblotting

Melanoma cells treated with ARN22089 for 6 or 24 h and lysed in RIPA buffer (EMD Millipore or an optimized cocktail (250 mM NaCL, 50 mM TrisCL pH 7.5, 0125% Nadeoxycholate, 0.375% Triton-X100, 0.15% NP-40, 4 mM EDTA) containing protease and phosphatase inhibitors cocktail (Thermo Scientific) and 1 mM PMSF and DTT. Lysates were then subjected to SDS-PAGE, transferred to PVDF membranes, followed by incubation with the indicated primary antibody and appropriate HRP-secondary antibody. ImageJ was used to perform densitometry.

A375 cells were maintained in DMEM (plus high glucose, L-glutamine, sodium pyruvate) supplemented with 10% FBS, 1% MEM NEAA and 1% antibiotic-antimycotic. WM3248 cells were maintained in 80% MCDB153 pH 7.30, 20% Leibovitz’s L-15, 2% FBS, 5 ug/mL insulin (bovine), 1.68 mM CaCl_2_, and 1% antibiotic-antimycotic. All cells kept in 5% CO_2_ incubator at 37°C.

### Reverse Phase Protein Arrays (RPPA)

Melanoma cells (WM3248) were seeded at 3.6 x 10^6^ cells per 150 mm plate and, 24 h later, treated with ARN22089 at 0, 5, 10, or 20 µM. Six hours later, cells were scraped and centrifuged at 1.3 KRPM for 15 min cold (between 4-16°C range). Pellets were snapped freeze in liquid nitrogen and shipped to MD Anderson Cancer Center core RPPA facility for RPPA analysis (**Table S5**). Three biological replicates for each sample were generated.

Multiple t test in Prism8 software was used to determine proteins with significant difference between 0 and 5, 10 or 20 µM. Normalized linear values were used to perform the t test with the following parameters: (1) Individual P values were computed with fewer assumptions by analyzing each row individually and did not assume consistent standard deviation; (2) Multiple comparisons were not corrected for; (3) alpha 0.05 was selected.

Heatmaps of RPPA represent normalized values (normalized linear values of 5, 10 or 20 µM divided by normalized linear values of 0 µM) (**Table S5-7**). Nomenclatures on the RPPA heatmaps: name_of_antibody.species_of_antibody.status_of_antibody. V - antibody validated; C - validation of antibody in progress; Q - antibody nonspecific; E - under evaluation; antibodies raised in M - mouse, G - goat, R - rabbit, or T - rat.

### RNA-seq analysis

WM3248 melanoma cells were seeded at 8.4 x 10^4^ cells. The next day, cells were treated with ARN22089 at 0, 4, or 6 µM and harvested 24 h later by adding RLT lysis buffer (Qiagen). Similarly, a small chunk of PDX tumor was collected and lysed in RLT lysis buffer. RNA was extracted using a RNeasy kit (Qiagen) and sent to the Genomic Core Facility at UCI for library construction and sequencing.

For the cell lines, paired-end sequencing reads were aligned to the human reference genome (GRCh37/hg19) with Tophat v2.1 (used in conjunction with Bowtie2 v2.2.7 and Samtools v1.9) and processed with Tuxedo Suite (Cufflinks v2.2.1).^63,64^ Heatmaps and other visualizations were generated using CummeRbund in R v4.0.3.

For PDX tumors, paired-end sequencing reads were aligned to both human and mouse reference genome (NCBI/GRCh38 and UCSC/mm10, respectively) using HISAT2 v2.1.0^65^; and Samtools v0.1.19 to convert to bam files and sort. Then XenofilteR^66^ was used to remove mouse reads, and reads were counted using featureCounts (subread v1.5.0-p3). Differential expression analysis was done using DESeq2^67^ in R.

Additional analyses of gene expressions were performed using STRING (‘Search Tool for Retrieval of Interacting Genes/Proteins’) and DAVID (‘Database for Annotation, Visualization hand Integrated Discovery’) to determine protein-protein network and functional annotation.^68,69^

### Bi-fluorescence Complementation Assay

Inducible HEK293 Flp-In T-REx cell line was a gift from Dr. Jean-Francois Cote in the Montreal Clinical Research Institute (IRCM), Montreal, Quebec, Canada.^70^ Cells were maintained in DMEM supplemented with 10% fetal bovine serum (TET tested, heat inactivated, R&D SYSTEMS INC), 1% antibiotic-antimycotic, 10 µg/mL blasticidin and 100 µg/mL zeocin at 37°C in 5% CO_2_. Stable BiFC lines were generated by co-transfecting pOG44 and pKK-BiFC-Venus- CDC42/RHOJ-PAK1 plasmids (9:1 ratio) and selecting with 100 µg/mL hygromycin and 10 µg/mL blasticidin 24-48 h after transfection. The CDC42 and RHOJ genes in the pKK-BiFC-Venus-CDC42/RHOJ-PAK1 plasmid, the following residues were modified to generate constitutively active form: CDC42, Q61L; RHOJ, Q79L.^71,72^

For CDC42 constitutively active (CA) assays, cells were seeded at 0.5 x 10^5^ cells per well of a 4-chamber slide. The next day, cells were treated with indicated doses and induced with 2 µg/mL doxycycline (DOX), simultaneously. Twenty-four hours later, cells were washed in PBS and fixed with 4% para-formaldehyde and 0.10% Triton-X100 in PBS for 15-20 min. Cells were then washed with PBS and incubated with blocking buffer (10% goat serum in PBS) for 30 min at RT and incubated with primary antibody (anti-GFP Invitrogen chicken Y fraction) at 1:1000 in blocking buffer 4 h RT, slow nutating. Cells were washed once with PBS buffer for 5 min. Secondary antibody (Alexa 488 nm) at 1:1000 in blocking buffer and cells were incubated for 1 h at RT, followed by DAPI 1:1000 in PBS for 20 min at RT. Cells were washed 3 times with PBS buffer, each time 5 min at RT. Slides were mounted with mounting medium (Vectorshield).

For RHOJ CA assays, cells were seeded at 1.5 x 10^5^ cells per well of a 4-chamber slide. Next day, cells were treated with indicated doses and induced with 2 µg/mL doxycycline (DOX), simultaneously. Eight hours later, cells were washed in PBS and fixed with 4% para-formaldehyde and 0.10% Triton-X100 in PBS for 15-20 min. Subsequent steps both clones were processed the same. Cells were then washed with PBS and incubated with blocking buffer (10% goat serum in 1 x PBS) for 30 min at RT and incubated with primary antibody (anti-GFP Invitrogen chicken Y fraction) at 1:2000 in blocking buffer overnight at 4°C, slow nutating. Cells were washed once with PBS buffer for 5 min. Secondary antibody (Alexa 488 nm) at 1:1000 in blocking buffer and cells were incubated for 1 h at RT, followed by DAPI 1:1000 in PBS for 20 min at RT. Cells were washed 3 times with PBS buffer, each time 5 min at RT. Slides were mounted with mounting medium (Vectorshield).

Slides were viewed using the Keyence BZ-X810 Wide-Field Microscope in the Stem Cell Research Center at the University of California, Irvine (UCI). Images were captured at high resolution with the same exposure time. Fluorescent images were quantified using Imaris software (BitPlane). Measurement setting includes surface masking [parameters: segment only a region of interest (excluding bright and blurry spots), classify surfaces, object-object statistics] to measure absolute intensity (threshold intensity adjusted for each image), and filter was set to “Number of Voxels Img=1” > 10. Intensity mean of all areas average from 5-6 images per condition at 20 x magnification. Percent average intensity mean was calculated as follow: ((Dose_1-NoDOX)/(DOX-NoDOX)) x 100). The standard error of mean was calculated taking the percent mean of intensity mean and divided by the square root of the number of surface areas measured.

### Vascularized Microtumor Assay

Methods for microfluidic device fabrication and establishing VMTs have been described previously (Sobrino 2016). Briefly, human endothelial colony-forming cell-derived endothelial cells (ECFC-EC) are isolated from cord blood via selection for the CD31+ cell population and cultured in EGM2 medium (Lonza). Normal human lung fibroblasts (LF) are purchased from Lonza. Cancer cells and LF are cultured in DMEM (Corning) containing 10% FBS (Gemini Bio). The ECFC-EC and cancer cells were transduced with lentivirus expressing mCherry (LeGO-C2, plasmid # 27339) or green fluorescent protein (GFP) (LeGO-V2, plasmid # 27340) (Addgene, Cambridge, Massachusetts). To load the microfluidic device, ECFC-EC and LF (both 8 x 10^6^ cells/mL) and cancer cells (2.5 × 10^6^ cells/mL) were resuspended in fibrinogen solution (10 mg/mL basal medium). The cell slurry was then mixed with 1 µL thrombin (3 U/mL) to catalyze gel solidification and quickly loaded into the tissue chambers of each VMT unit. Fibrin ECM was allowed to solidify at 37°C for 15 minutes prior to introducing ECM and EGM2 medium through the microfluidic channels. All cells and VMTs are cultured in a 37 °C, 20% O2, 5% O2 environment.

### Vascularized Microtumor Treatment

After culturing for 5 days to allow full development of each VMT, culture medium is replaced by medium containing the drugs (ARN22089, FRAX or vehicle) at the desired concentration and delivered through the microfluidic channels using the hydrostatic pressure gradient. Fluorescence images were acquired with an Olympus IX70 inverted microscope using SPOT software (SPOT Imaging, Sterling Heights, Michigan). AngioTool software (National Cancer Institute) was used to quantify vessel length in the VMT and ImageJ software (National Institutes of Health) was utilized to measure the total fluorescence intensity (i.e. mean grey value) for each tumor image to quantify tumor growth. Each chamber was normalized to time 0 baseline.

### Cancer cell line viability assay

Cancer cell line viability assay was performed by Proqinase with ARN22089 that we provided. Cells were cultured in their appropriate media and seeded in white multi-well plates. For the assays, cells were incubated with compound at 37°C overnight before compound was added. Cells were incubated in compound for 72 hours, and a CellTiterGlo assay was performed to quantify viable cells. The IC_50_ was calculated in a panel of 100 cell lines.

### PK Determination

ARN22089 was administered at 10 mg/kg PO and IP, while it was injected at 3 mg/kg IV to male mice. The vehicle for all administration routes was PEG400/Tween 80/Saline solution at 10/10/80% in volume, respectively. Three animals per dose/time point were treated, and blood samples were collected at selected time points up to 480 min. Plasma was separated from blood by centrifugation for 15 min at 1500 rpm a 4°C, collected in eppendorf tubes and frozen (−80°C). Control animals treated with vehicle only were also included in the experimental protocol. Plasma samples were centrifuged at 21.100 g for 15 min at 4°C. A 50 µl aliquot was transferred into a 96-Deep Well plate and 150 µl of extraction solution was added. The extraction solution was consisting of cold CH3CN spiked with 200 nM of internal standard. The plate was centrifuged at 21.100 g for 15 min at 4°C. Eight microliters of supernatant were then transferred into a 96-Deep Well plate and 80µl of H2O was added. A reference standard of the compound was spiked in naïve mouse plasma to prepare a calibration curve over a 1 nM – 10 µM range. Three quality control samples were prepared by spiking the compound in blank mouse plasma to the final concentrations of 20, 200 and 2000 nM. Calibrators and quality controls were extracted with the same extraction solution used for the plasma samples. The samples were analyzed on a Waters ACQUITY UPLC/MS TQD system (Waters Inc. Milford, USA) consisting of a TQD (Triple Quadrupole Detector) Mass Spectrometer equipped with an Electrospray Ionization interface and a Photodiode Array eλ Detector. The analyses were run on an ACQUITY UPLC BEH C18 (50 x 2.1 mmID, particle size 1.7 µm) with a VanGuard BEH C18 pre-column (5 x 2.1 mmID, particle size 1.7 µm) at 40°C. 0.1% HCOOH in H2O (A) and 0.1% HCOOH in CH3CN (B) were used as mobile phase with a linear gradient from 50 to 100% B in 2 min with the flow rate set to 0.5 mL/min. Electrospray ionization was applied in positive mode. Plasma levels of the parent compound was quantified by monitoring the MRM peak areas.

### Aqueous kinetic solubility assay

The aqueous kinetic solubility was determined from a 10mM DMSO stock solution of test compound in Phosphate Buffered Saline (PBS) at pH 7.4. The study was performed by incubation of an aliquot of 10mM DMSO stock solution in PBS (pH 7.4) at a target concentration of 250 μM resulting in a final concentration of 2.5% DMSO. The incubation was carried out under shaking at 25°C for 24 h followed by centrifugation at 14.800 rpm for 30 min. The supernatant was analyzed by UPLC/MS for the quantification of dissolved compound (in μM) by UV at a specific wavelength (215 nm). The UPLC/MS analyses were performed on a Waters ACQUITY UPLC/MS system consisting of a Single Quadrupole Detector (SQD) Mass Spectrometer (MS) equipped with an Electrospray Ionization (ESI) interface and a Photodiode Array Detector (PDA). The PDA range was 210-400 nm. ESI in positive mode was used in the mass scan range 100-650 Da. The analyses were run on an ACQUITY UPLC BEH C18 column (50 x2.1 mmID, particle size 1.7 µm) with a VanGuard BEH C18 pre-column (5 x 2.1 mmID, particle size 1.7 µm), using 10 mM NH_4_OAc in H_2_O at pH 5 adjusted with AcOH (A) and 10mM NH_4_OAc in CH_3_CN-H_2_O (95:5) at pH 5 (B) as mobile phase.

The thermodynamic solubility was determined by addition of Phosphate Buffered Saline (PBS) at pH 7.4 to an excess of solid test compound. The study was performed by incubation of an aliquot of 2.5mg of test compound in 500µL of PBS at pH 7.4 (final target concentration: 5mg/mL). The suspension was shaken at 300rpm for 24h at 25°C. The pH of the suspension was measured at the beginning and at the end of the incubation. At the end of the incubation, the saturated solution was filtered and analyzed by LC-MS for the quantification of dissolved compound by UV at 254nm using a calibration curve. The analyses were performed on a Waters ACQUITY UPLC-MS system consisting of a Single Quadrupole Detector (SQD) Mass Spectrometer equipped with an Electrospray Ionization interface and a Photodiode Array Detector from Waters Inc. (Milford, MA, USA). Electrospray ionization in positive mode was used in the mass scan range 100-650Da. The PDA range was 210-400nm. The analyses were run on an ACQUITY UPLC BEH C18 column (100×2.1mmID, particle size 1.7µm) with a VanGuard BEH C18 pre-column (5×2.1mmID, particle size 1.7µm), using 10mM NH_4_OAc in H2O at pH 5 adjusted with CH_3_COOH (A) and 10mM NH_4_OAc in CH3CN-H2O (95:5) at pH 5 (B) as mobile phase.

### *In vitro* mouse plasma stability assay

10mM DMSO stock solution of test compound was diluted 50-fold with DMSO-H2O (1:1) and incubated at 37°C for 2 h with mouse plasma added 5% DMSO (pre-heated at 37°C for 10 min). The final concentration was 2 µM. At each time point (0, 5, 15, 30, 60, 120 min), 50 µL of incubation mixture was diluted with 200 µL cold CH3CN spiked with 200 nM of internal standard, followed by centrifugation at 3.750 rpm for 20 min. The supernatant was further diluted with H2O (1:1) for analysis. The concentration of test compound was quantified by LC/MS-MS on a Waters ACQUITY UPLC/MS TQD system consisting of a Triple Quadrupole Detector (TQD) Mass Spectrometer (MS) equipped with an Electrospray Ionization (ESI) interface. The analyses were run on an ACQUITY UPLC BEH C18 column (50 x 2.1 mmID, particle size 1.7 µm) with a VanGuard BEH C18 pre-column (5 x 2.1 mmID, particle size 1.7µm) at 40°C, using H2O + 0.1% HCOOH in H2O (A) and CH3CN + 0.1% HCOOH (B) as mobile phase. ESI was applied in positive mode. The response factors, calculated on the basis of the internal standard peak area, were plotted over time. Response factor versus time profiles were fitted with Prism (GraphPad Software, Inc., USA) to estimate compounds half-life (t½) in mouse plasma.

### In vitro microsomal stability assay

10mM DMSO stock solution of each compound was pre-incubated at 37°C for 15 min with mouse liver microsomes in 0.1M Tris-HCl buffer (pH 7.4). The final concentration was 4.6μM. After pre-incubation, the co-factors (NADPH, G6P, G6PDH and MgCl_2_ pre-dissolved in 0.1M Tris-HCl) were added to the mixture and the incubation was continued at 37°C for 1h. At each time point (0, 5, 15, 30, 60min), 30μL of mixture was diluted with 200μL cold CH_3_CN spiked with 200nM of internal standard, followed by centrifugation at 3500g for 15min. The supernatant was then further diluted with H_2_O (1:1) for analysis. The concentration of each compound was quantified by LC/MS-MS on a Waters ACQUITY UPLC/MS TQD system consisting of a TQD (Triple Quadrupole Detector) Mass Spectrometer equipped with an Electrospray Ionization interface. The analyses were run on an ACQUITY UPLC BEH C18 (50×2.1mm ID, particle size 1.7μm) with a VanGuard BEH C18 pre-column (5×2.1mm ID, particle size 1.7μm) at 40°C, using 0.1% HCOOH in H_2_O (A) and 0.1% HCOOH in CH_3_CN (B) as mobile phases. The percentage of test compound remaining at each time point relative to the amount observed at t=0 was calculated. The half-lives (t½) were determined by a one-phase decay equation using a non-linear regression of compound concentration versus time.

### Mouse strains and PDX tumors

All animal experiments were approved by the UC Irvine Institutional Animal Care and Use Committee (IACUC) (AUP-17-230). NOD.Cg-*Prkdc^scid^ IL2rg^tm1Wjl^*/SzJ (NSG) mice were purchased from The Jackson Laboratories (stock number 005557). Human melanoma PDX tumors (563396-261-R, 128128-338-R, 174941-126-T, 156681-154-R, 425362-254-T) were purchased from Patient Derived Materials Repository (National Cancer Institute). Initial frozen tumors were divided into evenly sized pieces and kept viable in ice cold RPMI (Corning) with 10% FBS. Small incisions were made on both flanks of anesthetized NSG mice, with a single tumor piece placed under the skin near each incision. Tumors were allowed to form and reach 10% body mass before passaging into subsequent mice for drug studies. Excess portions of the tumor were frozen in RPMI with 10% FBS and 10% DMSO. Following tumor placement, mice were monitored until the tumors reached 100-200 mm^3^ to start drug treatment.

### Drug treatment for PDX tumors

ARN22089 was dissolved for 10 mg/kg doses in 100 µL of vehicle (80% Saline, 10% Tween80, and 10% PEG400). Stocks of concentrated drug were frozen at −20°C for long term and kept at 4°C for up to 5 days. Mice were treated with 100 µL of vehicle or RhoJ inhibitor twice daily for two weeks (morning and evening) through interperitoneal (IP) injections. Starting the same day as injections, tumor volumes were measured by caliper twice a week. Estimated tumor volumes were calculated by multiplying length by the width squared. At least four mice were imaged for each PDX during the course of treatment and continued until the tumors reached endpoint (1500 mm^3^). At endpoint, tumors were collected for histological analysis. For I.P. *BRAF^V600E^, PTPN11^N58S^* tumors; *BRAF^V600E^* – Vemurafenib Resistant tumors; *BRAF^V600E^, PTEN^H259Y^, TP53^C227Y^* tumors both male and female mice used in vehicle and ARN22089 treatment groups. All of the *BRAF^V600K^* tumors were grown in female mice, while all of the *BRAF^V600E^, PTEN^H259Y^, TP53^C227Y^* tumors were grown in male mice. Statistical analysis was completed on the tumor growth measurements using a two-way ANOVA comparing treatment groups for each PDX over time with TukeyHSD pairwise comparisons between timepoints in R (version 3.5.3).

Intravenous (IV) tail vein injections of ARN22089 were prepared at 10 mg/kg, 25 mg/kg, and 50 mg/kg as described above. Female mice were treated by IV twice a week as the *BRAF^V600E^, PTPN11^N58S^* tumors grew, for four weeks. Tumor and tissues were collected for histological analysis. Statistical analysis on tumor growth were completed in R (version 3.5.3) with a two-way ANOVA comparing treatment groups for each PDX over time with TukeyHSD pairwise comparisons between timepoints.

### Tissue Collection

Once tumors reached endpoint, various tissues were collected including the tumors, liver, and lymph nodes. All organs were fixed overnight in 10% formalin prior to washing and an ethanol dehydration series. Fixed tissue was sent to the Experimental Tissue Resource (UCI Department of Pathology) for embedding and preliminary staining by H&E and S100. The percentage of tumor necrosis was determined from H&E sections by a pathologist blinded to treatment groups. Statistical test on necrosis scores was completed using a student’s t-test in R (version 3.5.3).

## ACKNOWLEDGEMENTS

We thank Jessica Shiu, Zachary Springs, Pezhman Mobasher, Terry Nguyen, Celine Saade, Madeline McCanne, Katrina Huynh and and Giuliana Ottonello for their assistance with drug treatments and DMPK evaluation. We thank the co-founders of Alyra Therapeutics, Mark Benedyk and Alessandro Monge, for their advice and suggestions during the drug development and validation studies. This work was supported by grants from the National Cancer Institute (R01CA244571, U54CA217378) and the University of California, Irvine Chao Family Comprehensive Cancer Center Anti-Cancer Challenge to AKG and The Italian Foundation for Cancer Research (AIRC) (IG 18883 and IG 23679). JLF was supported by an NCI training grant to the Cancer Research Institute at the University of California, Irvine (T32CA009054-37).

The RPPA Core is supported by NCI Grant # CA16672 and Dr. Yiling Lu’s NIH R50 Grant # R50CA221675: Functional Proteomics by Reverse Phase Protein Array in Cancer. This work was made possible, in part, through access to the Genomics High Throughput Facility Shared Resource of the Cancer Center Support Grant (P30CA-062203) at the University of California, Irvine and NIH shared instrumentation grants 1S10RR025496-01, 1S10OD010794-01, and 1S10OD021718-01. The content is solely the responsibility of the authors and does not necessarily represent the official views of the National Institutes of Health.

## CONFLICT OF INTEREST

Anand Ganesan and Marco De Vivo are co-founders of a company entitled Alyra Therapeutics based on the technology presented in this manuscript.

