## Supplemental Figures for "Structure-based Design of CDC42 Effector Interaction Inhibitors For the Treatment of Cancer"

### Supplemental Figure 1

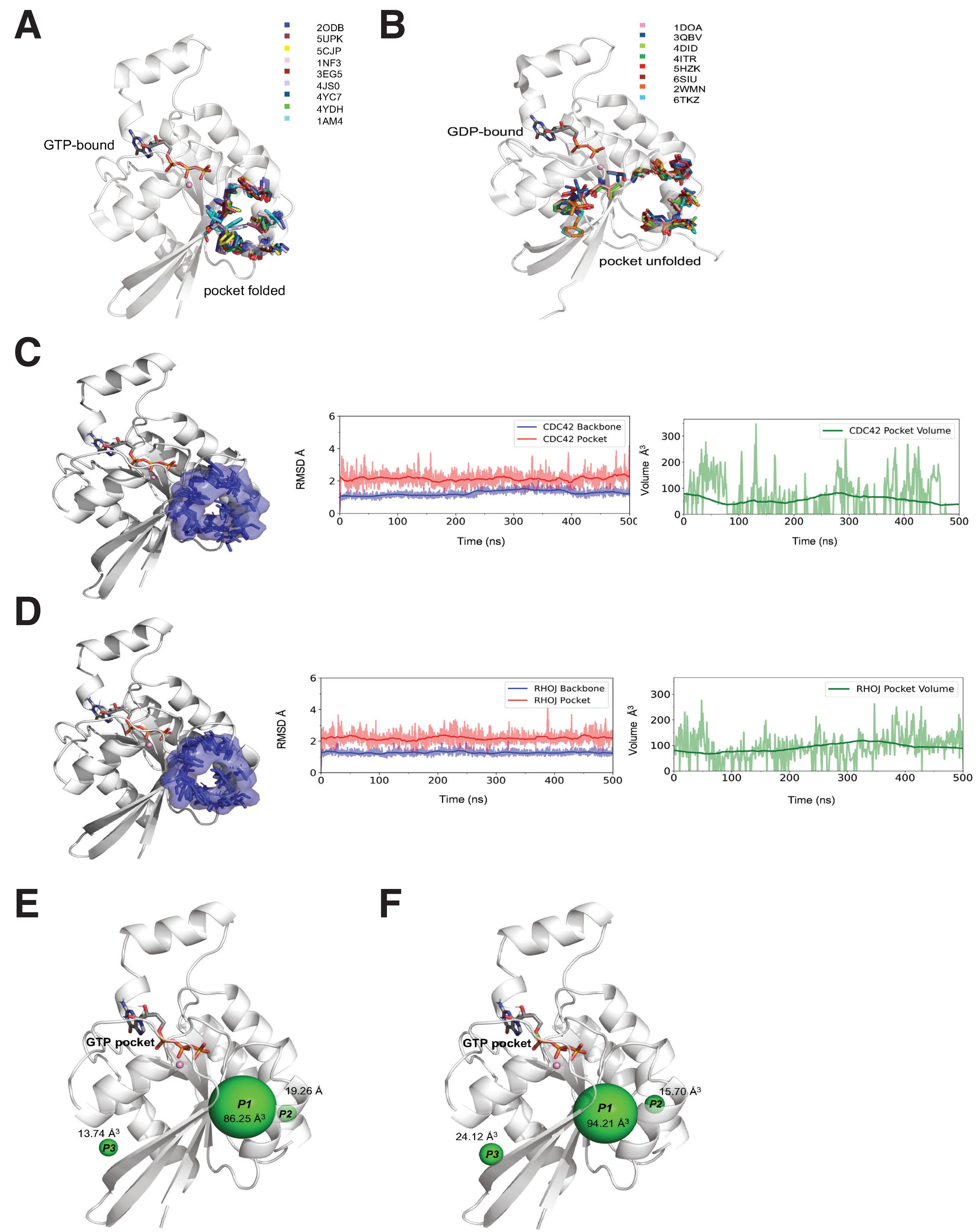

**Figure S1: Structural analysis of the CDC42 crystal structure** (see **Table S1**). Structures of GTP-bound CDC42 (**A**) and GDP-bound CDC42 (**B**) are reported. The protein is represented as cartoon, while the residues at the CDC42-effector interface are highlighted as sticks. Residue folds define the allosteric pocket only in the GTP-bound state (**A**). **MD simulations of GTP-bound CDC42 and RHOJ.** The structural representation of both GTP-bound CDC42 (**C**) and RHOJ (**D**) is reported on the left. Both CDC42 and RHOJ are represented as cartoon, while the binding pocket is highlighted as blue transparent surface. Multiple MD snapshots of the pocket residues (blue) are shown as sticks. On the right is reported the Root-Mean Square Deviation (RMSD) of the pocket and protein backbone and the volume of the pocket for both CDC42 (**C**) and RHOJ (**D**). The running average is in bold in each graph. **Binding pocket on CDC42 and RHOJ at the protein-protein interface, analyzed from MD simulations**. The structural representation of both GTP-bound CDC42 (**E**) and RHOJ (**F**) is reported, with the pocket represented in green spheres. The software pocketron was used to run the unsupervised analysis of our MD trajectories and identify such pockets at the protein-protein interface.[27] The volume of such pockets is reported for both the proteins.

### Supplemental Figure 2

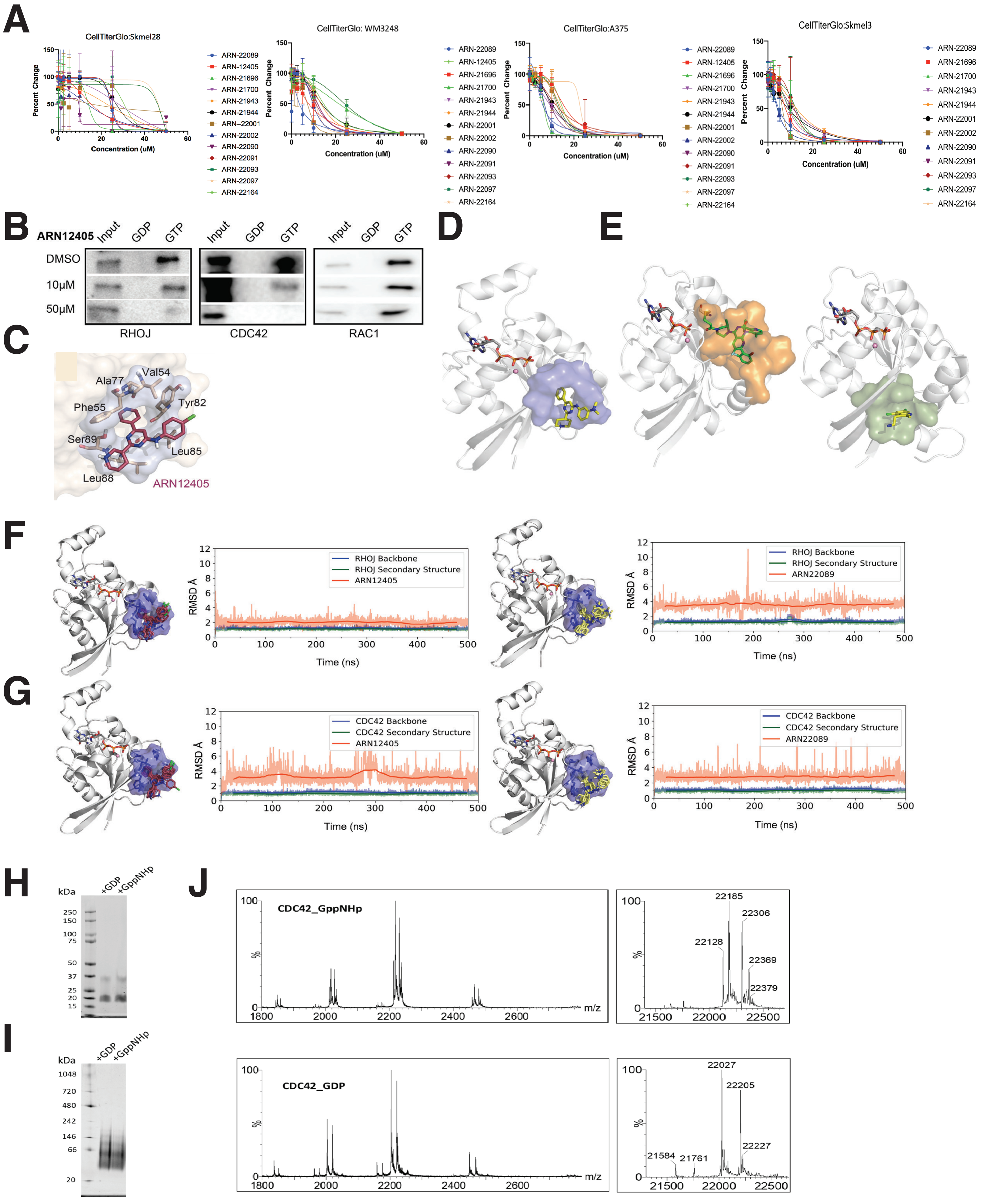

**Figure S2: Characterization of putative CDC42/RHOJ interaction inhibitors in multiple cell lines.** Graphical representation of the IC50 data displayed in Figure 2C. Results are from 3 independent experiments (five replicates per data point) in each of five cell lines **(A)**. **Lead compound inhibits RHOJ/CDC42 effector interactions**. WM3248 cells were treated with the indicated doses of ARN12405 and the effect on RHOJ/CDC42 interaction was measured using a CDC42 interaction assay **(B)**. Model structure of ARN12405 bound to CDC42/RHOJ is also shown (**C**). **Structural analysis and comparison of the pocket of the CDC42 GTPases compared to K-RAS**. (**D**) Structural representation of lead compound ARN22089 (yellow sticks) docked into the binding site (blue surface) of CDC42. **(E)** *Left*, The K-RAS-selective inhibitor DCAI bound to the pocket (green surface, PDB ID 4DST). *Right***,** The K-RAS(G12C) covalent inhibitors AMG-510 (green sticks, PDB ID 6OIM) at the inhibitor's pocket (orange surface). Notably, in (**E**), the pocket of (**D**) is missing. **MD simulations of protein-ligand complexes**. The structural representation of RHOJ (**F**) and CDC42 (**G**) in complex with either ARN12405 or ARN22089 is reported on the left. Both RHOJ and CDC42 are represented as cartoon, while the binding pocket is highlighted as blue transparent surface. Multiple MD snapshots of the ARN22089 (yellow) and ARN12405 (red) binding poses are shown as sticks. On the right, the RMSD over time for both RHOJ (**F**) and CDC42 (**G)** binding complexes. The RMSD running averages is in bold. **Quality control of His-CDC42 protein production and nucleotide loading**. GDP or GppNHp-loaded samples were compared of a His-CDC42 fragment (residues 4-182) using: **H)** SDS-PAGE 4-12 % v/v acrylamide; **I)** Native gel 4-16 % v/v acrylamide. **J)** ESI MS+ spectra and deconvoluted molecular weight of nucleotide bound His-CDC42. >98% of His-Cdc42 is estimated to be loaded with GppNHp and around 90% with GDP (with the remaining as apo-protein). Molecular weights + / - α-N-gluconoylation (+ 178 Da)[61]: 21584 / 21761 Da – apo-protein; 22027/ 22205 Da – His-Cdc42 + GDP; 22128 / 22306 Da – His-Cdc42 + GppNHp + $Mg^{2+}$; 22185 / 22369 Da – His-Cdc42 + GppNHp + $Mg^{2+}$ + $Ni^{2}$.

### Supplemental Figure 3

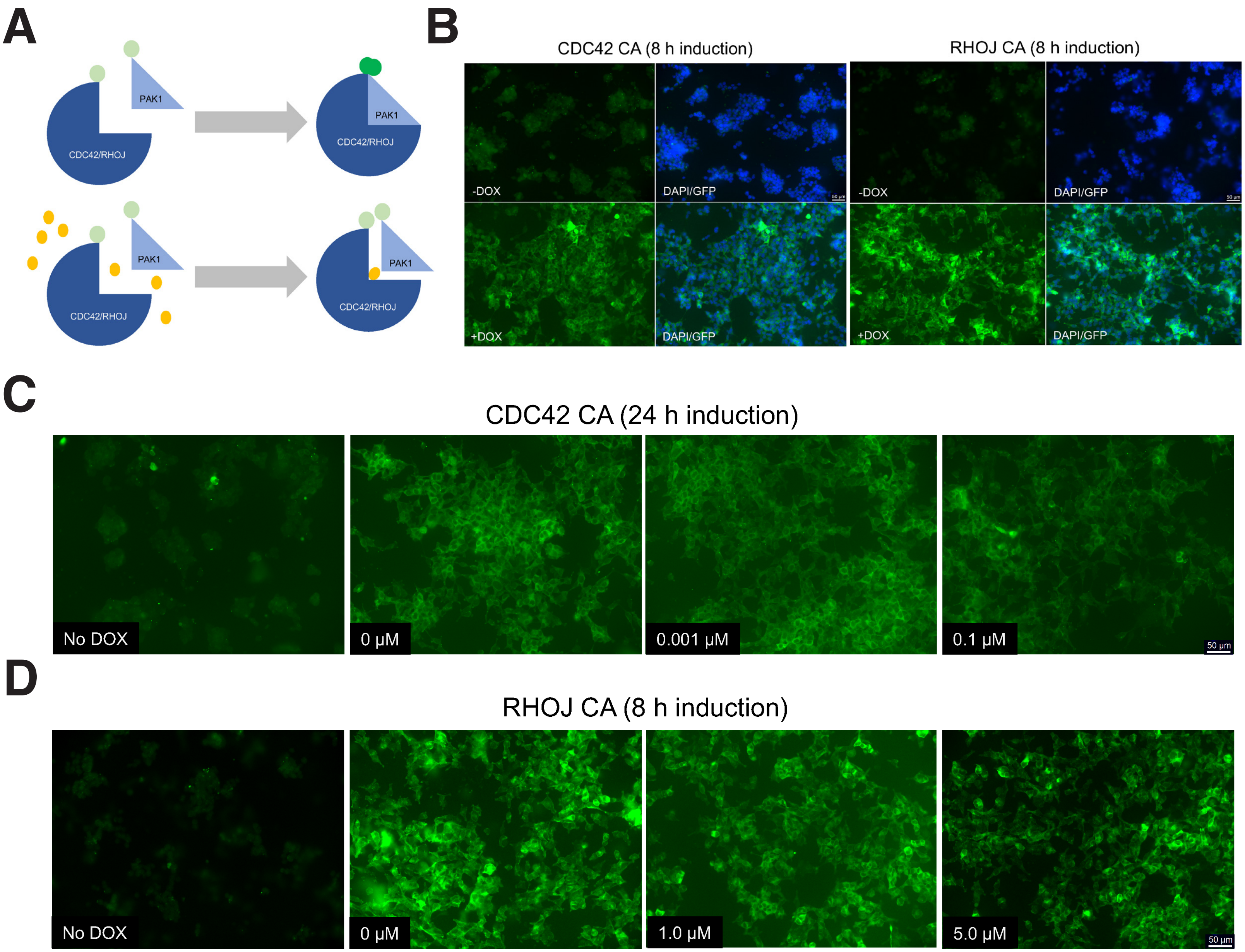

**Figure S3: Schematic diagram of the BiFC assay. (A)** In the absence of the lead drug, RHOJ/CDC42 and PAK1 bind and the cell exhibits YFP fluorescence. In the presence of the drug, RHOJ/CDC42 and PAK1 interaction is inhibited and no fluorescence is observed. **Development of a BiFC interaction assay to measure the cell activity of CDC42 interaction inhibitors.** Images of (**B**) CDC42 constitutive active (CA) and (**C**) RHOJ constitutive active (CA) BiFC cells. Cells were treated with or without DOX (2 μg/mL) for 8 h to induce protein expression. **CDC42 interaction inhibitors block CDC42 PAK interactions.** Representative images of BiFC cells induced (2 μg/mL DOX) and treated with indicated doses for (**D**) CDC42 CA at 0, 0.001, and 0.1 μM for 24 h and (**E**) RHOJ CA at 0, 1, 5 μM for 8 h.

### Supplemental Figure 4

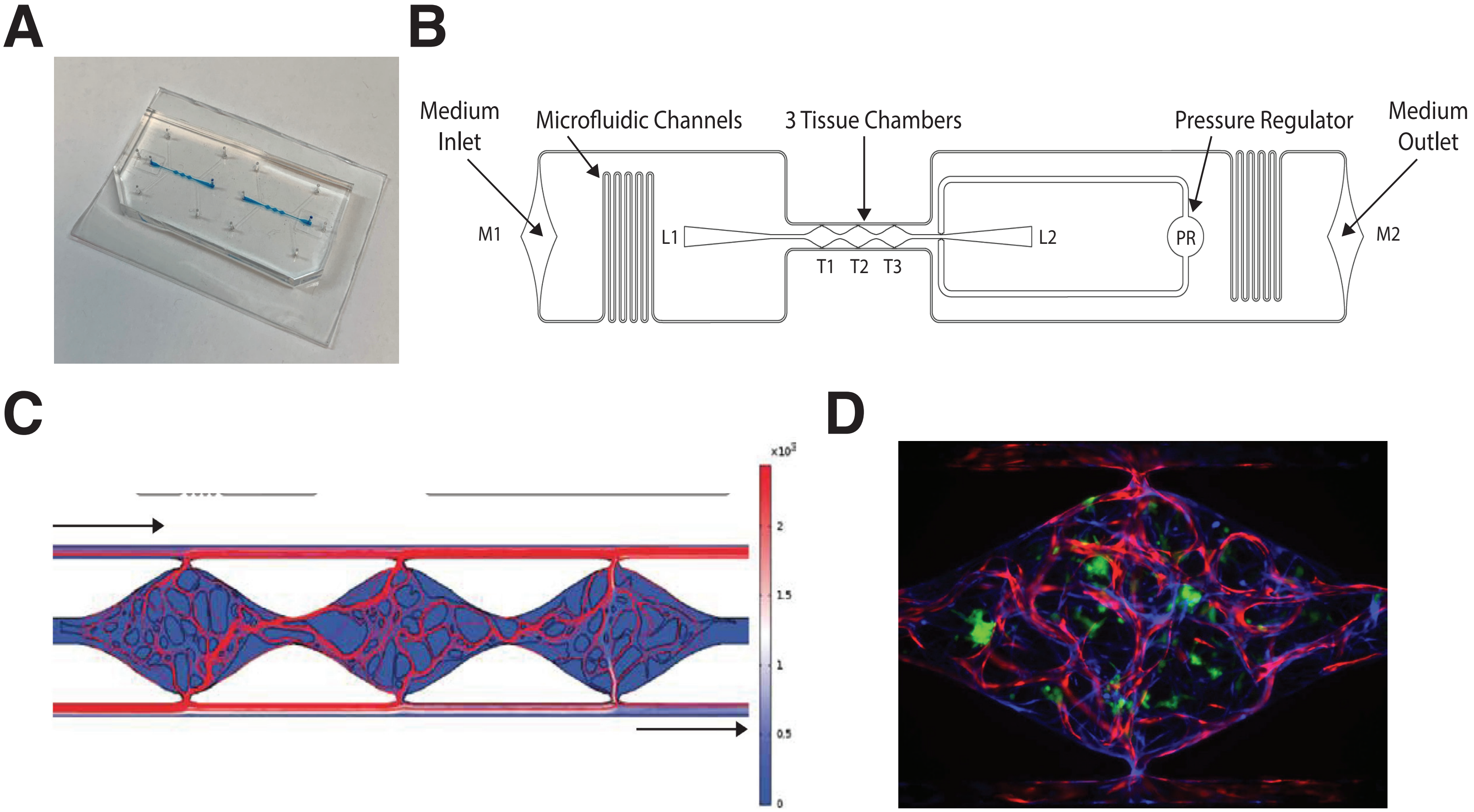

**Figure S4: Vascularized microtumor (VMT) model (see Figure 4). (A)** Photograph of microfluidic platform with blue dye injected into tissue chambers. (**B**) Zoom view shows a single device unit with 3 tissue chambers (T1-3) fed through microfluidic channels, 2 loading ports (L1-2), medium inlet and outlet (M1-2), and pressure regulator (PR) to prevent gel bursting during loading. (**C**) Simulation of flow through a formed and perfused vascular network. Top left is high pressure and bottom right is low pressure. (**D**) Fluorescent micrograph of the VMT with vessels labeled red (mCherry), fibroblasts labeled blue (azurite), and cancer cells labeled green (GFP). Chamber is 2 mm x 1 mm x 0.1 mm.

### Supplemental Figure 5

**A**

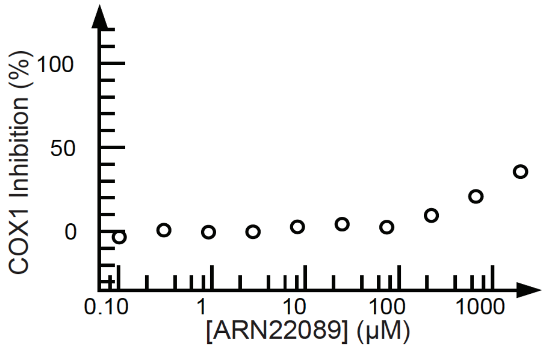


**B**

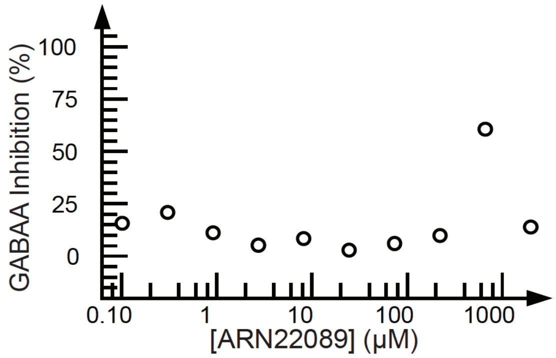


**C**

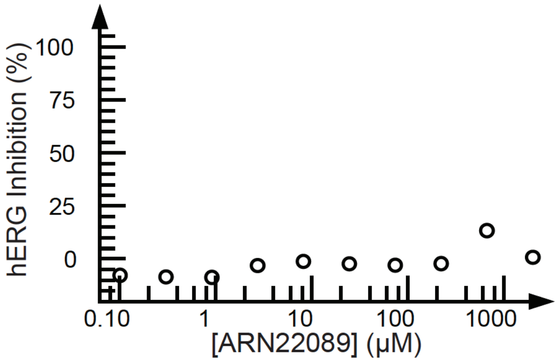


**D**

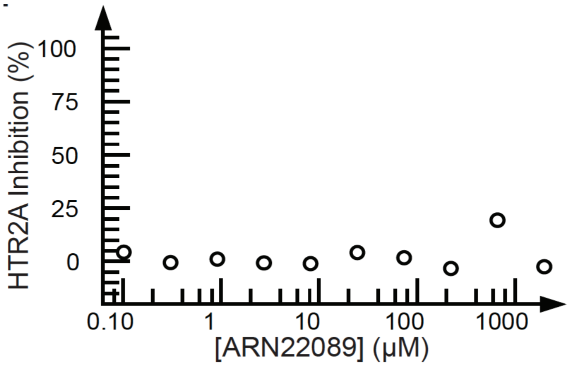


**E**

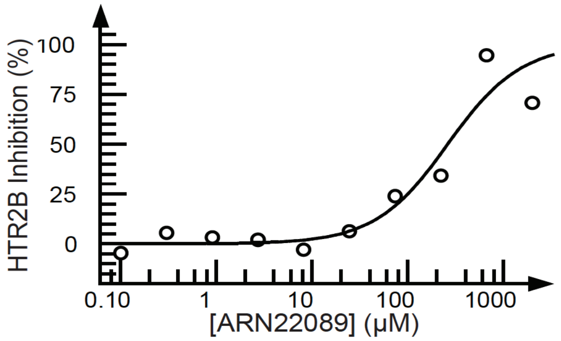


**F**

PK Parameters

| Parameter | Unit | I.P. (10mg/kg) | I.V. (3mg/kg) | P.O. (10mg/kg) |
|---|---|---|---|---|
| $C_{max}$ (obs) | ng/mL | 704 | 597 | 171 |
| $T_{max}$ (obs) | min | 15 | 5 | 120 |
| AUC (calc)[2] | ng-min/mL | 66707 | 63196 | 27780 |
| $T_{1/2}$ (calc) | min | 75 | 55 | 70 |
| $V_D$ (calc) | mL/kg | 14632 | 3627 | 30522 |
| CL (calc) | mL/min/kg | 135 | 46 | 303 |

**G**

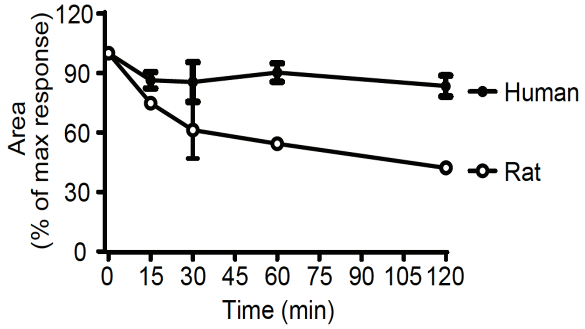


| | $T_{1/2}$ (min) | $CL_{INT}$ (ml/min/million cells) |
|---|---|---|
| Human | 510 | 0.002806 |
| Rat | 107 | 0.0271 |

**Figure S5: Activity of ARN22089 measured against a panel of common off-target proteins.** Curves for **(A)** COX1, **(B)** GABAA, **(C)** hERG, **(D)** HTR2A, **(E)** HTR2B are shown. **ARN22089 has drug like properties.** PK values of ARN22089 delivered I.P., I.V., and P.O. at the indicated doses was determined and is shown **(F)**. **ARN22089 is stable in plasma.** The stability of ARN22089 after incubation with human and rat liver microsomes was determined **(G)**.

### Supplemental Figure 6

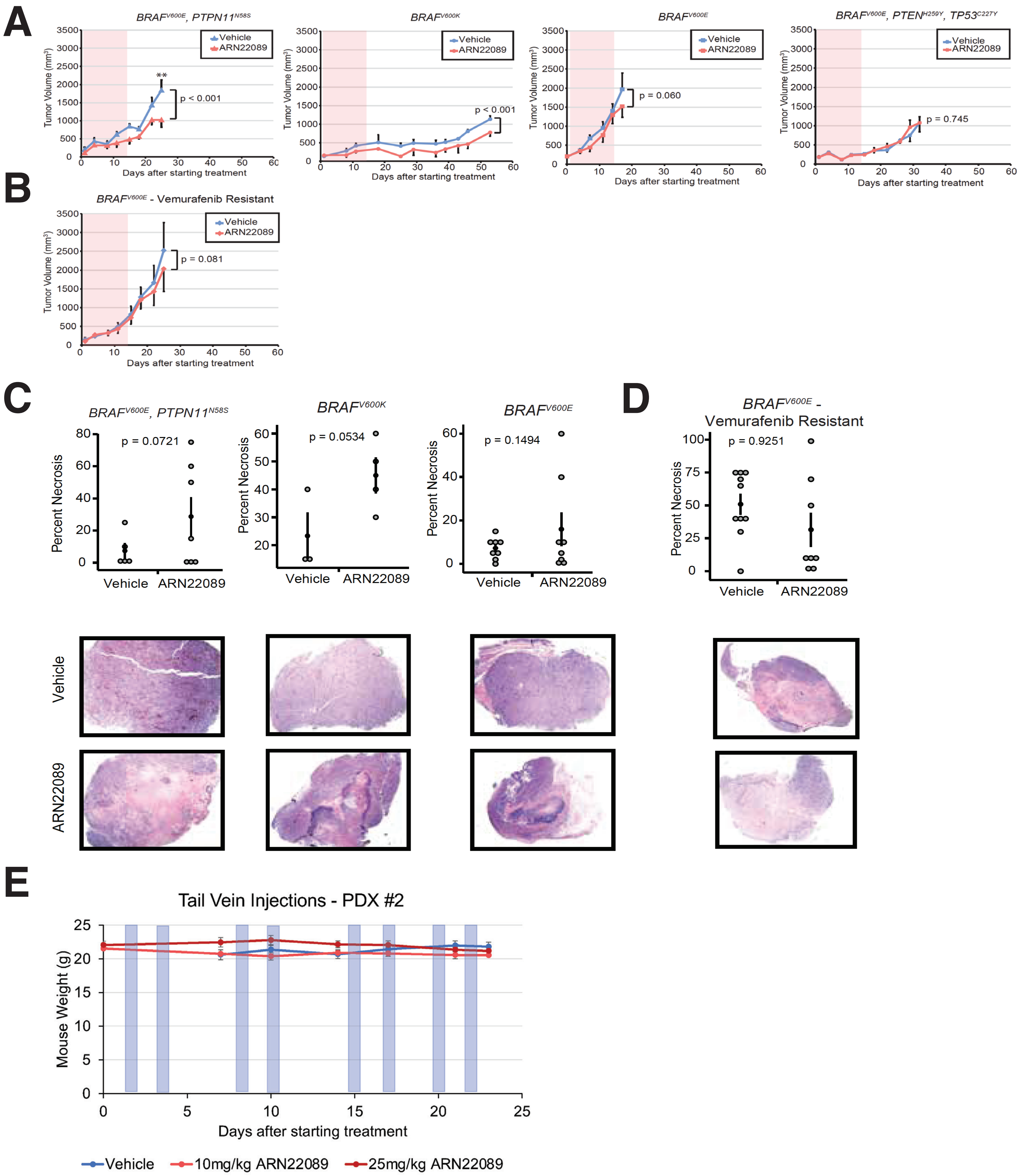

**Figure S6: ARN22089 inhibits tumor growth *in vivo*. Individual tumor growth curves for ARN22089 carrying PDX tumors.** Tumors with listed mutations treated with vehicle (blue) or 10 mg/kg ARN22089 (red) I.P. for 14 days. Treated days by I.P. are indicated by red shading. $BRAF^{V600E}$, $PTPN11^{N58S}$, n=6 tumors; $BRAF^{V600K}$, n=4 tumors; $BRAF^{V600E}$, n=8 tumors, $BRAF^{V600E}$, $PTEN^{H259Y}$, $TP53^{C227Y}$, n=6 vehicle and 8 ARN22089 treated tumors**.** Two-way ANOVA statistical tests compared treatment groups for each PDX over time with TukeyHSD pairwise comparisons between timepoints; * $p < 0.05$, ** $p < 0.01$, *** $p < 0.001$ (**A**). **BRAF inhibitor resistant tumors can respond to CDC42 inhibitors.** $BRAF^{V600E}$ – Vemurafenib Resistant, n=10 tumors were treated with ARN22089 as described in (**A**) and shown (**B**), **ARN22089 induces tumor necrosis.** Degree of necrosis present in hematoxylin and eosin (H&E) stained sections was determined by a blinded pathologist; n=5 vehicle and 7 ARN22089 treated for $BRAF^{V600E}$, $PTPN11^{N58S}$; n=3 vehicle and 4 ARN22089 treated for $BRAF^{V600k}$; n=7 vehicle and 8 ARN22089 treated $BRAF^{V600E}$;(shown in (**C**)) and n=10 vehicle and 8 ARN22089 treated for $BRAF^{V600E}$ – Vemurafenib Resistant (shown in (**D**) Necrosis scores for vehicle and ARN22089 treated tumors were compared using student's t-test; *$p < 0.05$. Representative images of H&E slides of vehicle and ARN22089 treated tumors shown below. **ARN22089 treatment does not induce cachexia**. Line graph represent mouse weights (g) during treatment, blue shaded boxes indicate I.V. treatments (see **Fig.6D**) (shown in (**E**)).

#### SUPPORTING INFORMATION


- **Title page and table of contents S1**
- **1. Chemistry: general experimental S2**
- **1.1. General Scheme of total synthesis of compounds ARN22097, ARN22093, ARN22091, ARN22164, ARN22090, ARN22089. S3**
- **1.2. Total synthesis of compound ARN22097 S3**
  **1.2.2. Representative Procedure 1 S4**
  **1.2.4. Representative Procedure 2 S5**
  **1.2.5. Representative Procedure 3 S6**
  **1.2.6. Representative Procedure 4 S6**
- **1.3. Total synthesis of compound ARN22093 S8**
- **1.4. Total synthesis of compound ARN22091 S10**
- **1.5. Total synthesis of compound ARN22164 S11**
- **1.6.Total synthesis of compound ARN22090 S13**
  **1.6.2. Representative Procedure 5 S15**
- **1.7. Total Synthesis of compound ARN22089 S16**
- **1.8. Binding evaluation of hit and derivatives to CDC42 by MST S17-S21**
- **1.9 Evaluation of protein:ARN22089 complex fluorescence change by SD-TEST S22**

#### 1. Chemistry: general experimental

Chemistry General considerations. All the commercial available reagents and solvents were used as purchased from vendors without further purification. Dry solvents were purchased from Sigma-Aldrich. Automated column chromatography purifications were done using a Teledyne ISCO apparatus (CombiFlash® Rf) with pre-packed silica gel columns of different sizes (from 4 g up to 24 g) and mixtures of increasing polarity of cyclohexane and ethyl acetate (EtOAc) or dichloromethane (DCM) and methanol (MeOH). NMR experiments were run on a Bruker Avance III 400 system (400.13 MHz for $^1H$, and 100.62 MHz for $^{13}C$), equipped with a BBI probe and Z-gradients. Spectra were acquired at 300 K, using deuterated dimethylsulfoxide (DMSO–$d_6$) or deuterated chloroform ($CDCl_3$) as solvents. For $^1H$ NMR, data are reported as follows: chemical shift, multiplicity (s= singlet, d= doublet, dd= double of doublets, t= triplet, q= quartet, p= pentuplet, m= multiplet), coupling constants (Hz) and integration. UPLC/MS analyses were run on a Waters ACQUITY UPLC/MS system consisting of a SQD (Single Quadrupole Detector) Mass Spectrometer equipped with an Electrospray Ionization interface and a Photodiode Array Detector. PDA range was 210-400 nm. The analyses were performed on an ACQUITY UPLC BEH C18 (50x2.1 mmID, particle size 1.7μm) with a VanGuard BEH C18 pre-column (5x2.1 mmID, particle size 1.7 μm) (LogD>1). The mobile phase was 10mM $NH_4OAc$ in $H_2O$ at pH 5 adjusted with AcOH (A) and 10mM $NH_4OAc$ in $CH_3CN$-$H_2O$ (95:5) at pH 5 (B). Electrospray ionization in positive and negative mode was applied in the mass scan range 100-500 Da. Depending on the analysis method used, a different gradient increasing the proportion of mobile phase B was applied. For analysis method A, the mobile-phase B proportion increased from 5 % to 95 % in 3 min. For analysis method B, the mobile-phase B proportion increased from 50 % to 100 % in 3 min. All final compounds displayed ≥ 93% purity as determined by UPLC/MS analysis. Compounds ARN22097, ARN22091, ARN22093, ARN22089, ARN22090, ARN22164 were synthesized in house, compounds ARN12405, ARN21698, ARN21700, 21699 were purchaised from Sigma-Aldrich and Asinex.

##### 1.1. General Scheme of total synthesis of compounds ARN22097, ARN22093, ARN22091, ARN22164, ARN22090, ARN22089.

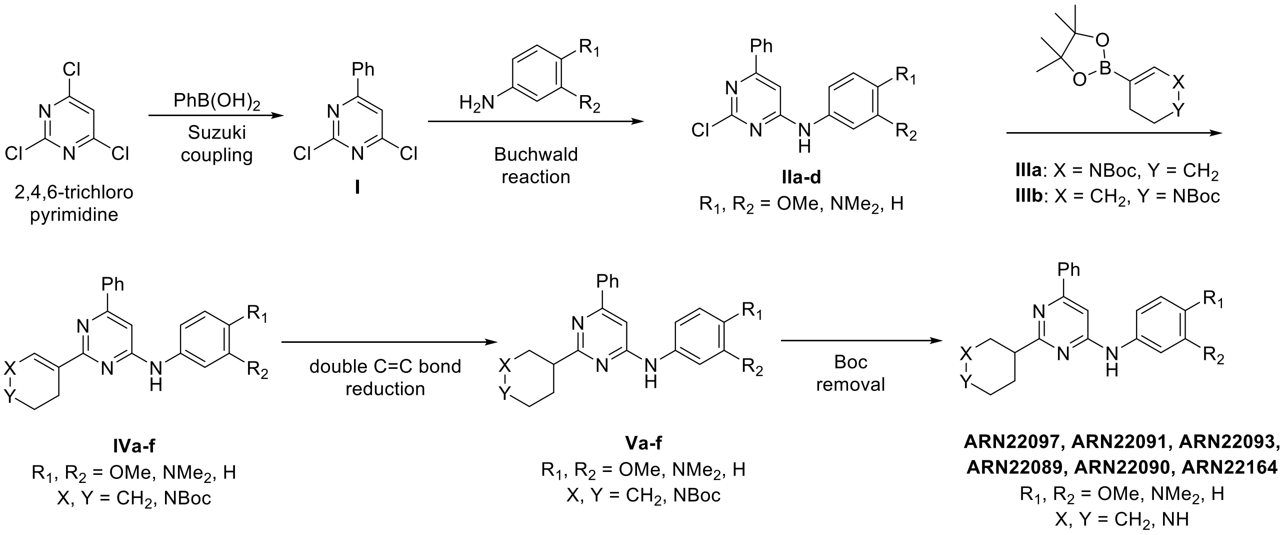


##### 1.2. Synthesis of compound ARN22097.

###### 1.2.1. Preparation of 2,4-dichloro-6-phenylpyrimidine (I).

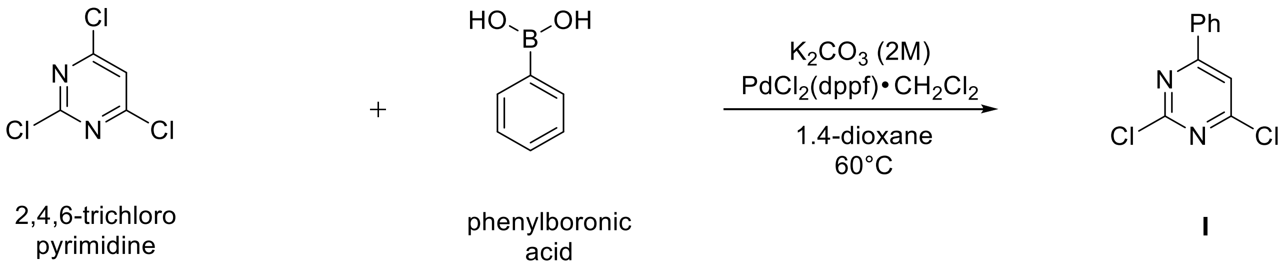


A suspension of 2,4,6-trichloropyrimidine (1000 mg, 5.29 mmol), phenyl boronic acid (665 mg, 5.29 mmol), dichloro[1,1'-bis(diphenylphosphino)ferrocene] palladium dichloromethane complex (204 mg, 0.26 mmol) and $K_2CO_3$ 2 M solution (5.3 ml, 10.58 mmol) in 1,4-dioxane (26.4 ml) was stirred in a CEM® microwave apparatus at 60 °C (200 W) for 1 hour. Resulting crude was portioned between dichloromethane (150 ml), $NaHCO_3$ saturated solution (100 ml), the organic layer dried over $Na_2SO_4$ and concentrated to dryness at low pressure. Final normal phase purification (cyclohexane/DCM from 100/0 to 85/15) afforded pure title compound (857 mg, yield 72 %). Rt = 1.38 min (analysis method 2); MS (ESI) m/z: 225.1 $[M-H]^+$, $[M-H]^+$ calculated: 225.0. $^1H$ NMR (400 MHz, $CDCl_3$) δ 8.13 – 8.03 (m, 2H), 7.68 (s, 1H), 7.62 – 7.48 (m, 3H).

###### 1.2.2. Representative procedure 1. Preparation of 2-chloro-*N*-(4-methoxyphenyl)-6-phenylpyrimidin-4-amine (IIa).

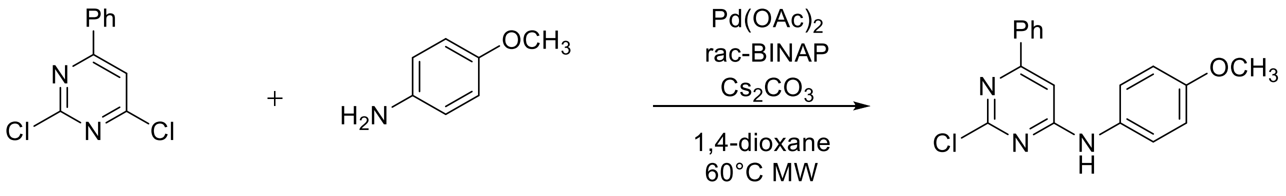


**I** 4-methoxyaniline **IIa**

A mixture of $Pd(OAc)_2$ (4.5 mg, 0.02 mmol, 0.05 equiv) and *rac*-BINAP (12.5 mg, 0.02 mmol, 0.05 equiv) in 1,4-dioxane (1.3 ml) was stirred under Ar flushing for 10 minutes. Then were stepwise added a solution of intermediate **I** (100 mg, 0.44 mmol, 1 equiv) in 1,4-dioxane (0.44 ml), a solution of 4-methoxyaniline (54.2 mg, 0.44 mmol, 1 equiv) in 1,4-dioxane (0.44 ml) and $Cs_2CO_3$ (172.0 mg, 0.52 mmol, 1.2 equiv). The reaction mixture was stirred in a CEM® microwave apparatus at 60 °C (200 W) for 4 hours, filtrated through a celite coarse patch, rinsed with DCM and concentrated to dryness at low pressure. Final normal phase purification (cyclohexane/TBME from 95/5 to 75/25) afforded pure title compound **IIa** (86.8 mg, yield 63 %). Rt = 1.39 min (analysis method 2); MS (ESI) m/z: 312.1 $[M\text{-}H]^+$, $[M\text{-}H]^+$ calculated: 312.1. $^1H$ NMR (400 MHz, DMSO-$d_6$) δ 9.91 (s, 1H), 8.06 – 7.84 (m, 2H), 7.65 – 7.37 (m, 5H), 7.06 (s, 1H), 7.02 – 6.92 (m, 2H), 3.76 (s, 3H).

**1.2.3. Preparation of tert-butyl 5-(4,4,5,5-tetramethyl-1,3,2-dioxaborolan-2-yl)-3,4-dihydropyridine-1(2H)-carboxylate (IIIa).**

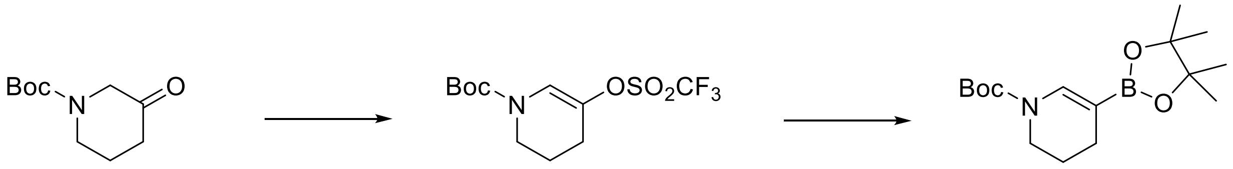


*tert*-butyl 3-oxopiperidine 1-carboxylate

*tert*-butyl 5-(((trifluoromethyl)sulfonyl)oxy) 3,4-dihydropyridine-1(2*H*)-carboxylate

**IIIa**

Step1: Preparation of tert-butyl 5-(((trifluoromethyl)sulfonyl)oxy)-3,4-dihydropyridine-1(2H)-carboxylate.

At -78 °C, to a solution of lithium diisopropylamide 2.0 M in cyclohexane (8.8 ml, 17.53 mmol) in dry tetrahydrofuran (15.6 ml) was added drop wise a solution of 3-oxo-piperidine-1- carboxyilic acid tert-butyl ester (3000 mg, 14.60 mmol) in dry tetrahydrofuran (15.6 mL). The mixture was stirred at -78 °C for 1 h and a solution of N-phenyl bis trifluoromethanesulfonamide (5855.6 mg, 16.07 mmol) in dry tetrahydrofuran (15.8 mL) was added. The mixture was stirred at -78 °C for 2 h and then was allowed to warm up to room temperature and stirred 16 additional hours at room temperature. The mixture was evaporated to dryness and the residue was taken with diethyl ether (50 ml), washed with water (50 mL), a 2 M solution of sodium hydroxide (50 mL) and brine (50 mL), dried over sodium sulfate and concentrated to dryness at low pressure. Final normal phase purification (cHexane/DCM from 100/0 to 50/50) afforded pure title compound (1354 mg, 28 % yield). Rt = 2.66 min (analysis method 1). $^1$H NMR (400 MHz, $CDCl_3$) δ 7.07 (s, 1H), 3.52 (s, 2H), 2.43 (td, $J$ = 6.4, 1.5 Hz, 2H), 1.93 (tt, $J$ = 6.3, 5.0 Hz, 2H), 1.49 (s, 9 H).

Step2: Preparation of tert-butyl 5-(4,4,5,5-tetramethyl-1,3,2-dioxaborolan-2-yl)-3,4-dihydropyridine-1(2H)-carboxylate (**IIIa**).

To a degassed solution of sulfonate derived from step 1 (600 mg, 1.81 mmol) in 1,4-dioxane (10.7 ml) was added bis-(pinacolato)diboron (603.8 mg, 2.35 mmol), potassium acetate (502.7 mg, 5.07 mmol) and dichloro[1,1'-bis(diphenylphosphino)ferrocene] palladium dichloromethane complex (139.5 mg, 0.18 mmol) were added. The mixture was stirred at 80 °C for 3 h. After cooling down, the mixture was filtered and resulting filtrate concentrated to dryness at low pressure. Final normal phase purification (cHexane/DCM from 70/30 to 50/50) afforded pure title compound **IIIa** (448 mg, 80 % yield). Rt = 1.85 min (analysis method 2). MS (ESI) m/z 310.2 $[M-H]^+$, $[M-H]^+$ calculated: 310.2. $^1$H NMR (400 MHz, $CDCl_3$) δ 5.29 (s, 1H), 3.67 – 3.39 (m, 2H), 2.15 – 1.96 (m, 2H), 1.81 – 1.73 (m, 2H), 1.49 (s, 9H), 1.32 – 1.17 (m, 12H).

##### 1.2.4. Representative procedure 2. Preparation of tert-butyl 5-(4-((4-methoxyphenyl)amino)-6-phenylpyrimidin-2-yl)-3,4-dihydropyridine-1(2H)-carboxylate (IVa).

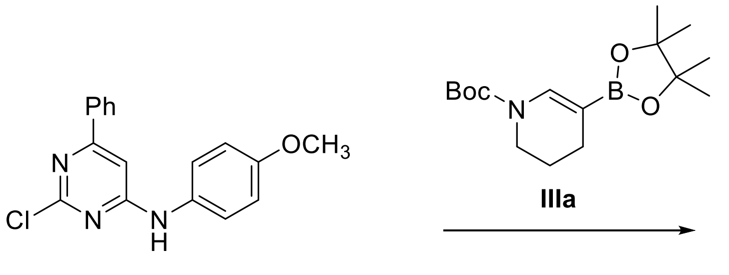


A suspension of intermediate **IIa** (100 mg, 0.32 mmol, 1 equiv), 4,4,5,5-tetramethylboronate **IIIa** (119.0 mg, 0.38 mmol, 1.2 equiv dichloro[1,1'-bis(diphenylphosphino)ferrocene] palladium dichloromethane complex (26.1 mg, 0.03 mmol, 0.1 equiv) and $K_2CO_3$ 2M solution (0.32 mL, 0.64 mmol, 2 equiv) in 1,4-dioxane (10 ml) was stirred in a CEM® microwave apparatus at 120 °C (200 W) for 2 hours. Resulting crude was portioned between dichloromethane (25 ml), $NaHCO_3$ saturated solution (25 ml), the organic layer dried over $Na_2SO_4$ and concentrated to dryness at low pressure. Final normal phase purification (cyclohexane/AcOEt from 100/0 to 80/20) afforded pure title compound **IVa** (64.7 mg, yield 44 %). Rt = 2.37 min (analysis method 2); MS (ESI) m/z 459.6 $[M-H]^+$, $[M-H]^+$ calculated: 459.2. $^1$H NMR (400 MHz, $CDCl_3$) δ 8.05 – 7.97 (m, 2H), 7.46 – 7.37 (m, 3H), 7.31 (t, $J$ = 6.5 Hz, 2H), 6.98 – 6.90 (m, 2H), 6.71 (s, 1H), 6.68 – 6.46 (m, 1H), 3.84 (s, 3H), 3.66 (d, $J$ = 9.5 Hz, 2H), 2.82 – 2.60 (m, 2H), 2.02 – 1.86 (m, 2H), 1.56 (s, 9H).

**1.2.5. <u>Representative procedure 3.</u> Preparation of tert-butyl 3-(4-((4-methoxyphenyl)amino)-6-phenylpyrimidin-2-yl)piperidine-1-carboxylate (Va).**

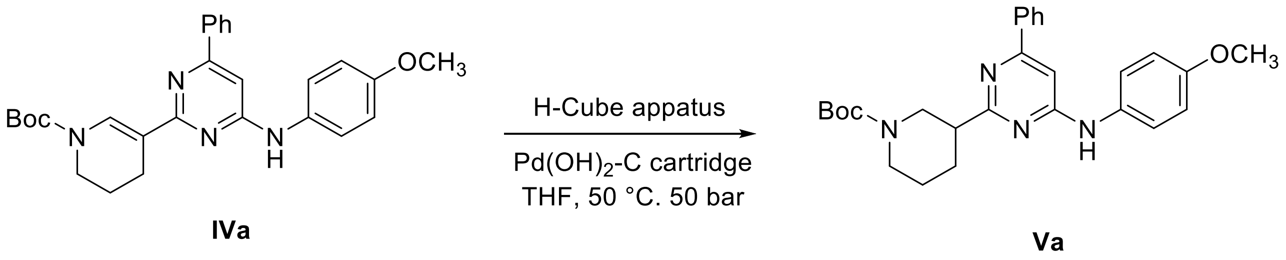


A solution of intermediate **IVa** (65.0 mg, 0.14 mmol) in THF (1.4 mL) was eluted in a H-cube apparatus through a $Pd(OH)_2$-C cartridge at 50 °C under 50 bar $H_2$ pressure until reaction completion. Final normal phase purification (cHexane/TBME from 100/0 to 80/20) afforded pure title compound **Va** (40 mg, yield 62 %). Rt = 1.94 min (analysis method 2); MS (ESI) m/z: 461.2 $[M-H]^+$, $[M-H]^+$ calculated: 461.2. $^1H$ NMR (400 MHz, $CDCl_3$) δ 8.07 – 7.80 (m, 2H), 7.46 – 7.38 (m, 3H), 7.26 (s, 1H), 7.01 – 6.89 (m, 2H), 6.77 (s, 1H), 6.71 (s, 1H), 3.84 (s, 3H), 3.01 – 2.68 (m, 2H), 2.23 (d, $J$ = 12.3 Hz, 1H), 1.80 (qd, $J$ = 13.0, 3.8 Hz, 2H), 1.73 – 1.52 (m, 4H), 1.47 (s, 9H).

**1.2.6. <u>Representative procedure 4</u>. Preparation of N-(4-methoxyphenyl)-6-phenyl-2-(piperidin-3-yl)pyrimidin-4-amine (ARN22097).**

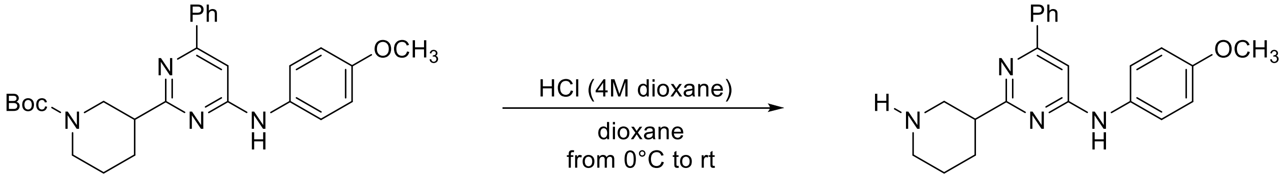


**VIa** **ARN22097**

To a 0 °C solution of protected intermediate **VIa** (65.0 mg, 0.14 mmol, 1 equiv) in 1,4-dioxane (0.36 ml) was dropwise added a HCl (4M) solution in 1,4-dioxane (0.35 mL, 1.4 mmol, 10 equiv) and the reaction mixture stirred at room temperature for 1 h, then the reaction crude was concentrated to dryness at low pressure, the resulting crude portioned between DCM (4 mL) and NaOH 0.1 M (4 mL), the organic layer dried over $Na_2SO_4$ and concentrated to dryness at low pressure. Final normal phase purification (elution by gradient from 85/15 to 65:35 DCM/DCM:$NH_3$ 1M MeOH 4:1) afforded pure title compound **ARN22097** (31 mg, yield 99 %). Rt = 0.49 min (analysis method 2); MS (ESI) m/z: 361.6 $[M-H]^+$, $[M-H]^+$ calculated: 361.2. $^1H$ NMR (400 MHz, DMSO-$d_6$) δ 9.60 (s, 1H), 8.15 – 7.90 (m, 2H), 7.59 (d, $J$ = 8.4 Hz, 2H), 7.56 – 7.47 (m, 3H), 7.05 (d, $J$ = 4.2 Hz, 1H), 7.00 – 6.90 (m, 2H), 3.76 (s, 3H), 3.59 (dd, $J$ = 18.0, 10.2 Hz, 1H), 3.27 – 3.18 (m, 3H), 2.94 – 2.87 (m, 1H), 2.25 – 2.19 (m, 1H), 1.91 – 1.83 (m, 2H), 1.79-1.69 (m, 1H). $^{13}C$ NMR (100 MHz, DMSO-$d_6$) δ 168.5 (Cq), 161.3 (Cq), 161.0 (Cq), 155.0 (Cq), 137.1

(Cq), 132.8 (Cq), 130.2 (CH), 128.8 (CH, 2C), 126.4 (CH, 2C), 121.7 (CH, 2C), 114.0 (CH, 2C), 99.4 (CH), 55.2 ($CH_3$), 46.1 ($CH_2$), 43.1 ($CH_2$), 41.1 (CH), 27.9 ($CH_2$), 21.5 ($CH_2$).

#### 1.3. Synthesis of compound ARN22093.

##### 1.3.1. Preparation of *N*1-(2-chloro-6-phenylpyrimidin-4-yl)-*N*4,*N*4-dimethylbenzene-1,4-diamine (IIb).

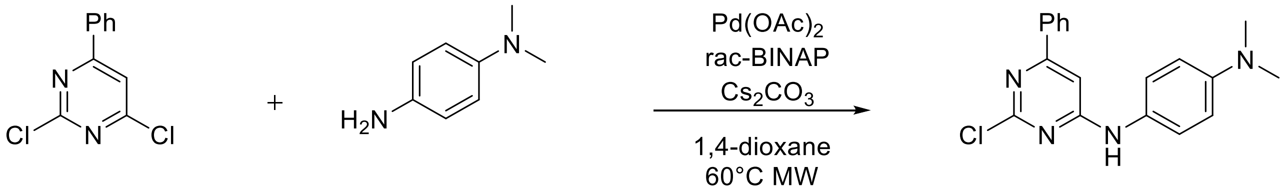


**I** **IIb** **IIIb**

Compound **IIIb** was prepared according Representative Procedure 1: using intermediate **I** (300 mg, 1.33 mmol) and *N*1,*N*1-dimethylbenzene-1,4-diamine **IIb** (191.1 mg, 1.33 mmol). Final normal phase purification (cyclohexane/TBME from 100/0 to 80/20) afforded pure title compound **IIIb** (272 mg, yield 63 %). Rt = 1.56 min (analysis method 2); MS (ESI) m/z: 325.1 $[M-H]^+$, $[M-H]^+$ calculated: 325.1. $^1H$ NMR (400 MHz, DMSO-$d_6$) δ 9.76 (s, 1H), 7.93 (dd, J = 6.7, 3.0 Hz, 2H), 7.52 (dd, J = 4.6, 2.4 Hz, 3H), 7.36 (s, 2H), 7.00 (s, 1H), 6.87 – 6.64 (m, 2H), 2.89 (s, 6H).

##### 1.3.2. Preparation of tert-butyl 5-(4-((4-(dimethylamino)phenyl)amino)-6-phenylpyrimidin-2-yl)-3,4-dihydropyridine-1(2H)-carboxylate (IVb).

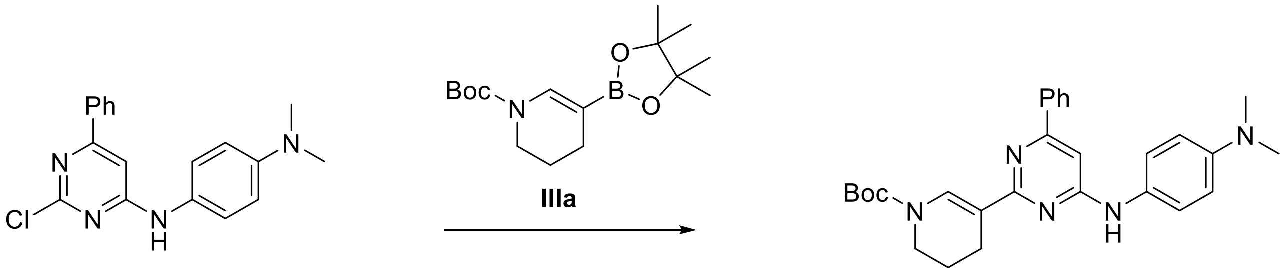


**IIb** **IVb**

Compound **IVb** was prepared according Representative Procedure 2: using intermediates **IIb** (100 mg, 0.35 mmol) and **IIIa** (117.8 mg, 0.37 mmol). Final normal phase purification (cyclohexane/AcOEt from 100/0 to 85/15) afforded pure title compound **IVb** (107.4 mg, yield 74 %). Rt = 2.58 min (analysis method 2); MS (ESI) m/z 472.4 $[M-H]^+$, $[M-H]^+$ calculated: 472.3. $^1H$ NMR (400 MHz, DMSO-$d_6$) δ 9.28 (s, 1H), 8.12 – 7.96 (m, 2H), 7.64 – 7.42 (m, 5H), 7.15 (s, 1H), 6.96 (s, 1H), 6.86 – 6.69 (m, 2H), 4.11 (d, $J$ = 4.0 Hz, 2H), 3.55 (t, $J$ = 5.7 Hz, 2H), 2.87 (s, 6H), 2.67 (d, $J$ = 7.0 Hz, 2H), 1.44 (s, 9H).

##### 1.3.5. Preparation of tert-butyl 3-(4-((4-(dimethylamino)phenyl)amino)-6-phenylpyrimidin-2-yl)piperidine-1-carboxylate (Vb).

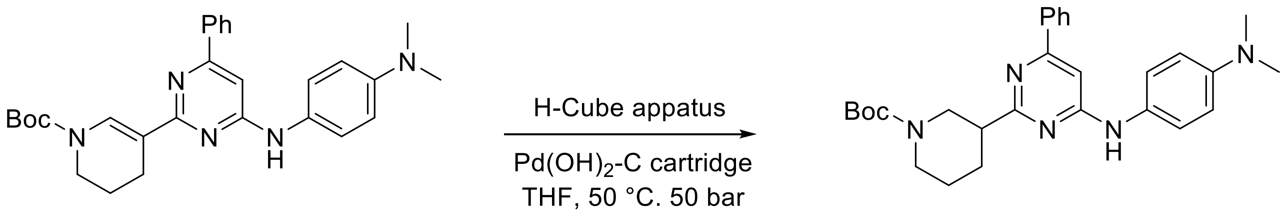


**IVb** **Vb**

Compound **Vb** was prepared according Representative Procedure 3: using intermediate **IVb** (60.5 mg, 0.12 mmol). Final normal phase purification (cHexane/TBME from 90/10 to 70:30) afforded pure title compound **Vb** (60 mg, yield 99 %). Rt = 2.22 min (analysis method 2); MS (ESI) m/z: 474.4 $[M-H]^+$, $[M-H]^+$ calculated: 474.3. $^1$H NMR (400 MHz, DMSO-$d_6$) δ 9.38 (s, 1H), 8.24 – 7.80 (m, 2H), 7.59 – 7.34 (m, 5H), 6.98 (s, 1H), 6.91 – 6.59 (m, 2H), 3.37 (d, $J$ = 9.8 Hz, 1H), 3.16 – 3.02 (m, 1H), 3.02 – 2.91 (m, 2H), 2.87 (s, 6H), 2.68 (td, $J$ = 11.9, 3.0 Hz, 1H), 2.23 – 2.05 (m, 1H), 1.89 – 1.70 (m, 2H), 1.63 (q, $J$ = 12.8 Hz, 1H), 1.45 (s, 9H).

##### 1.3.6. Preparation of *N1*,*N1*-dimethyl-*N4*-(6-phenyl-2-(piperidin-3-yl)pyrimidin-4-yl)benzene-1,4-diamine (ARN22093).

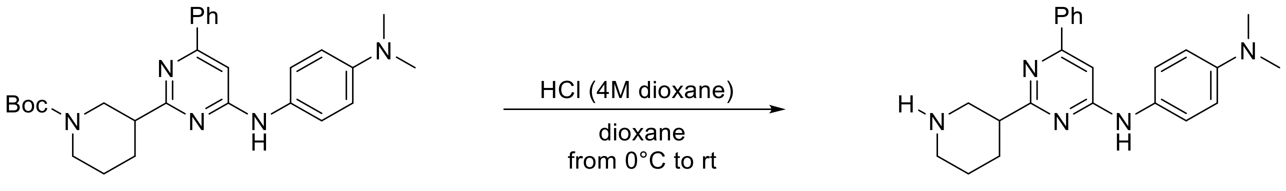


**VIb** **ARN22093**

Compound **ARN22093** was prepared according Representative Procedure 4: using intermediate **VIb** (61 mg, 0.13 mmol), and HCl (4M) solution in 1,4-dioxane (0.35 ml) in 1,4-dioxane (0.36 mL). Final normal phase purification (elution by gradient from 80/20 to 60/40 DCM/DCM:$NH_3$ 1M MeOH 4:1) afforded pure title compound **ARN22093** (46 mg, yield 95 %). UPLC_MS: Rt = 1.96 min (analysis method 1); MS (ESI) m/z: 374.6 $[M-H]^+$, $[M-H]^+$ calculated: 374.2. $^1$H NMR (400 MHz, DMSO-$d_6$) δ 9.38 (s, 1H), 7.99 – 7.96 (m, 2H), 7.52 – 7.44 (m, 5H), 6.98 (s, 1H), 6.67 – 6.33 (m, 2H), 3.37 (d, $J$ = 9.8 Hz, 1H), 3.16 – 3.02 (m, 1H), 3.00 – 2.90 (m, 2H), 2.87 (s, 6H), 2.68 (td, $J$ = 12.1, 3.0 Hz, 1H), 2.16 – 2.11 (m, 1H), 1.89 – 1.58 (m, 3H). $^{13}$C NMR (100 MHz, DMSO-$d_6$) δ 170.2 (Cq), 161.4 (Cq), 160.8 (Cq), 146.8 (Cq), 137.4 (Cq), 136.9 (Cq), 129.9 (CH), 128.7 (CH, 2C), 126.3 (CH, 2C), 121.9 (CH, 2C), 112.9 (CH, 2C), 98.6 (CH), 49.0 ($CH_2$), 44.9 ($CH_2$), 44.0 (CH), 40.5 ($CH_3$, 2C), 29.0 ($CH_2$), 24.1 ($CH_2$).

#### 1.4. Total synthesis of compound ARN22091.

##### 1.4.1. Preparation of 2-chloro-*N*-(3-methoxyphenyl)-6-phenylpyrimidin-4-amine (IIc).

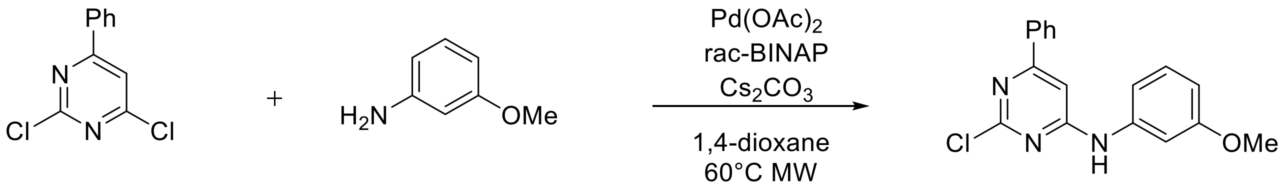


**I** 3-methoxyaniline **IIc**

Compound **IIc** was prepared according Representative Procedure 1: using intermediate **I** (300 mg, 1.13 mmol) and 3-methoxyaniline (154 µl, 1.33 mmol). Final normal phase purification (cyclohexane/TBME from 100/0 to 85/15) afforded pure title compound **IIc** (150 mg, yield 36 %). Rt = 1.52 min (analysis method 2); MS (ESI) m/z: 312.1 $[M-H]^+$, $[M-H]^+$ calculated: 312.1. $^1$H NMR (400 MHz, DMSO-$d_6$) δ 10.07 (s, 1H), 8.00 – 7.90 (m, 2H), 7.59 – 7.50 (m, 3H), 7.34 (t, $J$ = 2.3 Hz, 1H), 7.29 (t, $J$ = 8.1 Hz, 1H), 7.18 (s, 2H), 6.69 (ddd, $J$ = 8.2, 2.5, 0.9 Hz, 1H), 3.77 (s, 3H).

##### 1.4.2. Preparation of tert-butyl 5-(4-((3-methoxyphenyl)amino)-6-phenylpyrimidin-2-yl)-3,4-dihydropyridine-1(2H)-carboxylate (IVc).

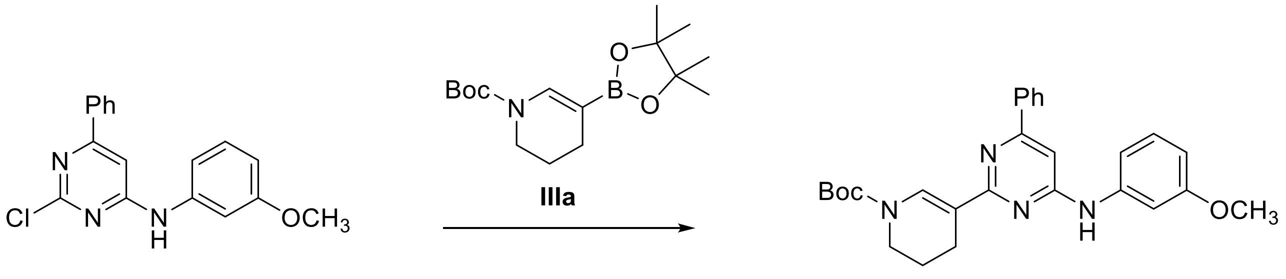


**IIc** **IVc**

Compound **IVc** was prepared according Representative Procedure 2: using intermediate **IIc** (100 mg, 0.32 mmol) and intermediate **IIIa** (119.0 mg, 0.38 mmol) following the general procedure reaction C previously described. Final normal phase purification (cyclohexane/TBME from 100/0 to 80/20) afforded pure title compound **IVc** (65.0 mg, yield 44 %). Rt = 2.40 min (method 2); MS (ESI) m/z 459.6 [M-H]+, [M-H]+ calculated: 459.2.

##### 1.4.3. Preparation of tert-butyl 3-(4-((3-methoxyphenyl)amino)-6-phenylpyrimidin-2-yl)piperidine-1-carboxylate (Vc).

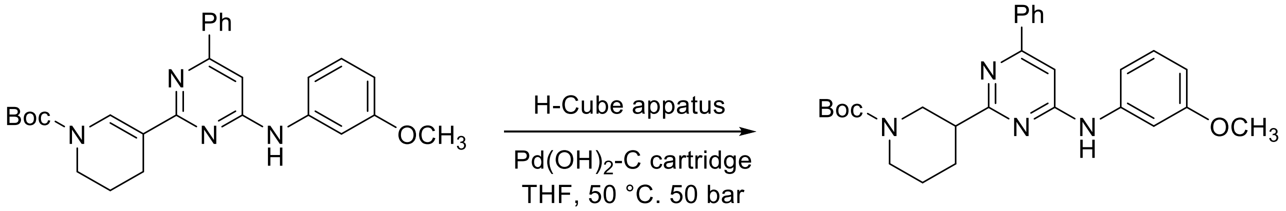


**IVc** **Vc**

Compound **Vc** was prepared according Representative Procedure 3: using intermediate **IVc** (65.0 mg, 0.14 mmol). Final normal phase purification (cHexane/AcOEt from 100/0 to 80:20) afforded pure title compound (32.6 mg, yield

50 %). Rt = 2.13 min (analysis method 2); MS (ESI) m/z: 461.6 [M-H]+, [M-H]+ calculated: 461.2. $^{1}$H NMR (400 MHz, $CDCl_3$) δ 8.04 – 7.92 (m, 2H), 7.45 (p, *J* = 3.9, 3.2 Hz, 3H), 7.30 (t, *J* = 8.1 Hz, 1H), 7.08 (t, *J* = 2.2 Hz, 1H), 6.99 (s, 1H), 6.95 (dd, *J* = 7.9, 2.0 Hz, 1H), 6.89 (s, 1H), 6.72 (dd, *J* = 8.3, 2.4 Hz, 1H), 3.84 (s, 3H), 3.20 (s, 1H), 2.99 – 2.88 (m, 1H), 2.80 (t, *J* = 12.5 Hz, 1H), 2.26 (d, *J* = 12.5 Hz, 1H), 1.93 – 1.74 (m, 2H), 1.75 – 1.51 (m, 3H), 1.47 (s, 9H).

##### 1.4.4. Preparation of *N*-(3-methoxyphenyl)-6-phenyl-2-(piperidin-3-yl)pyrimidin-4-amine (ARN22091).

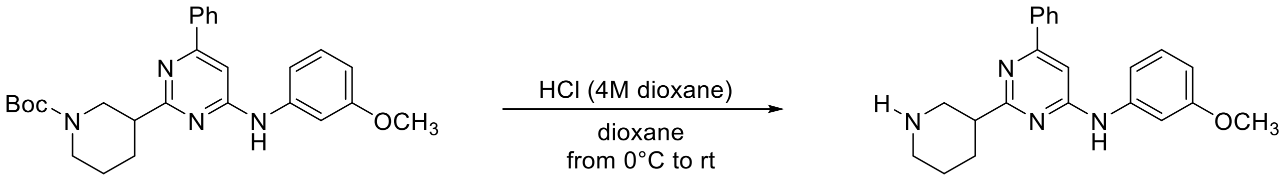


**VIc** **ARN22091**

Compound **ARN22091** was prepared according Representative Procedure 4: using intermediate **VIc**, (61 mg, 0.13 mmol), and HCl (4M) solution in 1,4-dioxane (0.35 ml) in 1,4- (0.36). Final normal phase purification (elution by gradient from 85/15 to 60/40 DCM/DCM:$NH_3$ 1M MeOH 4:1) afforded pure title compound 7 (39 mg, yield 82 %). UPLC_MS: Rt = 1.91 min (analysis method 1); MS (ESI) m/z: 361.6 $[M-H]^{+}$, $[M-H]^{+}$ calculated: 361.2. $^{1}$H NMR (400 MHz, DMSO-$d_6$) 9.89 (s, 1H), 8.17 – 7.94 (m, 2H), 7.67 – 7.44 (m, 4H), 7.34 – 7.12 (m, 3H), 6.61 (dt, *J* = 5.4, 2.4 Hz, 1H), 3.79 (s, 3H), 3.63 (d, *J* = 8.1 Hz, 1H), 3.27 – 3.18 (m, 3H), 2.89 (d, *J* = 14.1 Hz, 1H), 2.26 (d, *J* = 11.7 Hz, 1H), 2.01 – 1.67 (m, 3H). $^{13}$C NMR (100 MHz, DMSO-$d_6$) δ 168.5 (Cq), 161.3 (Cq), 161.1 (Cq), 159.6 (Cq), 141.2 (Cq), 136.9 (Cq), 130.3 (CH), 129.4 (CH), 128.8 (CH, 2C), 126.4 (CH, 2C), 111.8 (CH), 108.0 (CH), 105.1 (CH), 100.5 (CH), 55.0 ($CH_3$), 46.1 ($CH_2$), 43.1 ($CH_2$), 41.2 (CH), 27.9 ($CH_2$), 21.5 ($CH_2$).

#### 1.5. Total synthesis of compound ARN22164.

##### 1.5.1. Preparation of *N1*-(2-chloro-6-phenylpyrimidin-4-yl)-*N3,N3*-dimethylbenzene-1,3-diamine

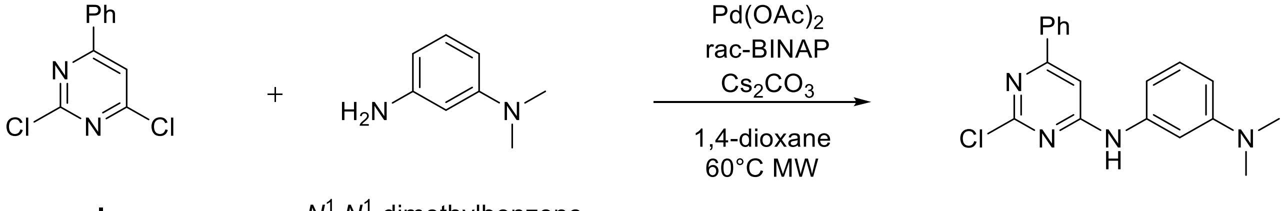


**I** $N^1$,$N^1$-dimethylbenzene-1,3-diamine **IId**

Compound **IId** was prepared according Representative Procedure 1: using intermediate **I** (300 mg, 1.13 mmol) and N1,N1-dimethylbenzene-1,3-diamine (181.5 mg, 1.33 mmol). Final normal phase purification (cyclohexane/TBME from 100/0 to 80/20) afforded pure title compound (246 mg, yield 57 %). Rt = 1.73 min (analysis method 2); MS (ESI) m/z: 325.1 $[M-H]^{+}$, $[M-H]^{+}$ calculated: 325.1. $^{1}$H NMR (400 MHz, DMSO-$d_6$) δ 9.93 (s, 1H), 8.14 – 7.82 (m,

2H), 7.60 – 7.47 (m, 3H), 7.29 – 7.09 (m, 2H), 7.04 (s, 1H), 6.99 – 6.85 (m, 1H), 6.51 (ddd, *J* = 8.4, 2.5, 0.8 Hz, 1H), 2.92 (s, 6H).

**1.5.2. Preparation of tert-butyl 5-(4-((3-(dimethylamino)phenyl)amino)-6-phenylpyrimidin-2-yl)-3,4-dihydropyridine-1(2H)-carboxylate (IVd).**

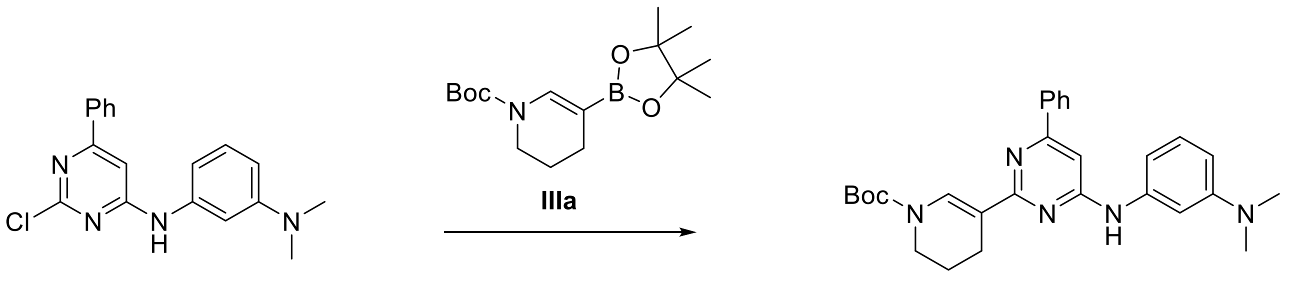


**IId** **IVd**

Compound **IVd** was prepared according Representative Procedure 2: using **IId** (175 mg, 0.54 mmol) and intermediate **IIIa** (199.9 mg, 0.65 mmol). Final normal phase purification (cyclohexane/AcOEt from 100/0 to 85/15) afforded pure title compound (109.2 mg, yield 43 %). Rt = 2.65 min (analysis method 2); MS (ESI) m/z 472.3 [M-H]$^+$, [M-H]$^+$ calculated: 472.3. $^1$H NMR (400 MHz, DMSO-$d_6$) δ 9.34 (s, 1H), 8.40 (s, 1H), 8.04 (dd, *J* = 7.7, 1.9 Hz, 2H), 7.59 – 7.44 (m, 4H), 7.11 (t, *J* = 8.0 Hz, 2H), 6.99 (s, 1H), 6.40 (dd, *J* = 9.1, 2.5 Hz, 1H), 3.59 (t, *J* = 5.6 Hz, 2H), 2.92 (s, 6H), 2.65 – 2.59 (m, 2H), 1.88 (p, *J* = 6.0 Hz, 2H), 1.50 (s, 9H).

**1.5.3. Preparation of tert-butyl 3-(4-((3-(dimethylamino)phenyl)amino)-6-phenylpyrimidin-2-yl)piperidine-1-carboxylate (Vd).**

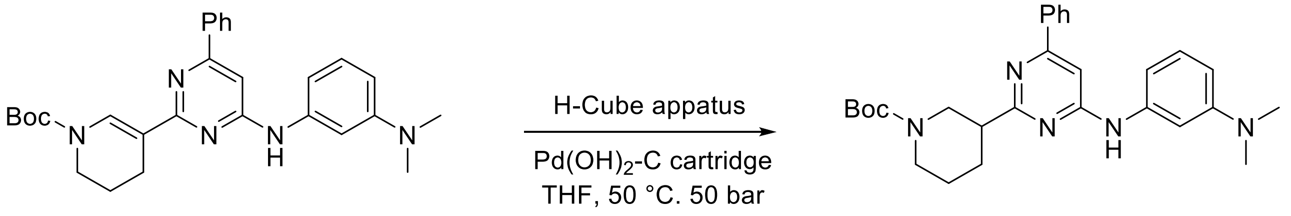


**IVd** **Vd**

Compound **Vd** was prepared according Representative Procedure 3: using intermediate **IVd** (105 mg, 0.22 mmol). Final normal phase purification (cHexane/AcOEt from 100/0 to 80/20) afforded pure title compound Vd (12 mg, yield 12 %). Rt = 2.35 min (analysis method 2); MS (ESI) m/z: 474.6 [M-H]$^+$, [M-H]$^+$ calculated: 474.3. $^1$H NMR (400 MHz, $CDCl_3$) δ 8.04 – 7.93 (m, 2H), 7.49 – 7.37 (m, 4H), 7.04 (s, 1H), 6.78 (s, 1H), 6.69 (d, *J* = 7.7 Hz, 1H), 6.59 (d, *J* = 8.5 Hz, 1H), 4.20 – 4.08 (m, 1H), 3.23 – 3.16 (m, 1H), 2.99 (s, 6 H), 2.98 – 2.91 (m, 1H), 2.88 – 2.77 (m, *J* = 14.3 Hz, 1H), 2.31 – 2.20 (m, 1H), 1.82 – 1.58 (m, 2H), 1.51 – 1.45 (m, 11H).

**1.5.4. Preparation of *N1*,*N1*-dimethyl-*N3*-(6-phenyl-2-(piperidin-3-yl)pyrimidin-4-yl)benzene-1,3-diamine (ARN22164)**

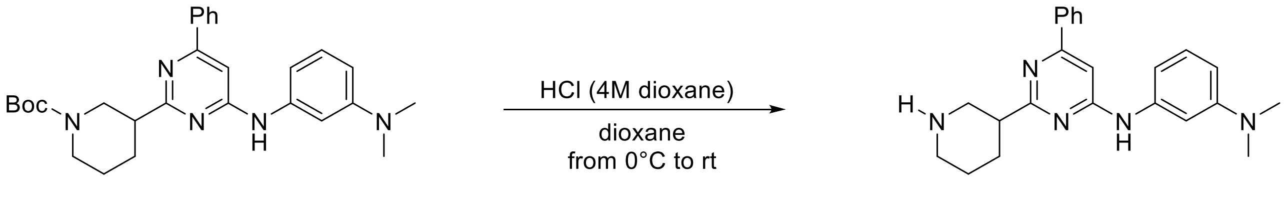


**VId** **ARN22164**

Compound **ARN22164** was prepared according Representative Procedure 4: using intermediate **VId**, (34 mg, 0.08 mmol), and HCl (4M) solution in 1,4-dioxane (0.22 ml) in 1,4-dioxane (0.22). Final normal phase purification (elution by gradient from from 95/5 to 45/55 DCM/DCM:$NH_3$ 1M MeOH 4:1) afforded pure title compound **ARN22164** (16 mg, yield 61 %). UPLC_MS: Rt = 2.07 min (analysis method 1); MS (ESI) m/z: 374.5 $[M-H]^+$, $[M-H]^+$ calculated: 374.2. $^1H$ NMR (400 MHz, DMSO-$d_6$) δ 9.50 (s, 1H), 8.01 (dd, $J$ = 7.8, 1.8 Hz, 2H), 7.56 – 7.48 (m, 3H), 7.45 (s, 1H), 7.12 (t, $J$ = 8.1 Hz, 1H), 7.10 – 7.06 (m, 1H), 6.90 (d, $J$ = 8.0 Hz, 1H), 6.40 (dd, $J$ = 8.2, 2.5 Hz, 1H), 3.26 – 3.17 (m, 1H), 2.94 (s, 6H), 2.94 – 2.87 (m, 1H), 2.82– 2.75 (m, 2H), 2.46 (dd, $J$ = 12.1, 2.9 Hz, 1H), 2.16 – 2.03 (m, 1H), 1.90 – 1.72 (m, 1H), 1.71 – 1.58 (m, 1H), 1.57 – 1.39 (m, 1H). $^{13}C$ NMR (100 MHz, DMSO-$d_6$) δ 171.5 (Cq), 161.3 (Cq), 160.9 (Cq), 150.9 (Cq), 141.0 (Cq), 137.4 (Cq), 130.0 (CH), 129.0 (CH), 128.8 (CH, 2C), 126.4 (CH, 2C), 107.8 (CH), 106.7 (CH), 103.8 (CH), 99.7 (CH), 51.4 ($CH_2$), 46.4 (CH), 46.3 ($CH_2$), 40.3 ($CH_3$, 2C), 30.0 ($CH_2$), 26.1 ($CH_2$).

##### 1.6. Total synthesis of compound ARN22090.

###### 1.6.1. Preparation of tert-butyl 4-(4-((3-methoxyphenyl)amino)-6-phenylpyrimidin-2-yl)-3,6-dihydropyridine-1(2H)-carboxylate (IVe).

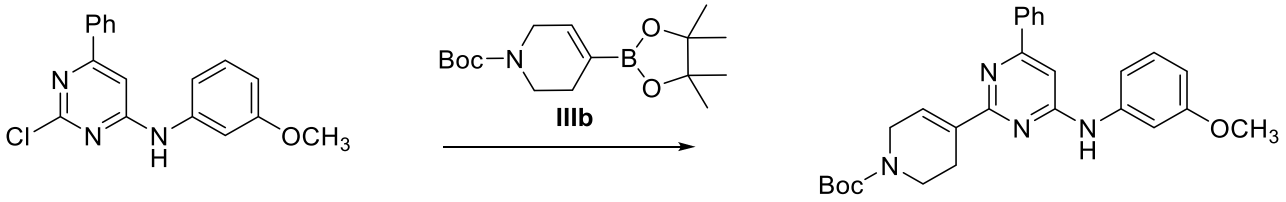


**IIc** **IVe**

Compound **IVe** was prepared according Representative Procedure 2: using intermediate **IIc** (180 mg, 0.58 mmol) and tert-butyl 4-(4,4,5,5-tetramethyl-1,3,2-dioxaborolan-2-yl)-3,6-dihydro-2H-pyridine-1-carboxylate **IIIb** (202.4 mg, 0.64 mmol). Final normal phase purification (cyclohexane/AcOEt from 100/0 to 85/15) afforded title compound **ARN22090** (241.2 mg, yield 91 %). Rt = 2.31 min (analysis method 2); MS (ESI) m/z 459.3 $[M-H]^+$, $[M-H]^+$ calculated: 459.2. $^1H$ NMR (400 MHz, DMSO-$d_6$) δ 9.63 (s, 1H), 8.19 – 8.03 (m, 2H), 7.62 – 7.46 (m, 4H), 7.33 – 7.22 (m, 2H), 7.20 (s, 1H), 7.11 (s, 1H), 6.67 – 6.54 (m, 1H), 4.13 (d, $J$ = 3.3 Hz, 2H), 3.79 (s, 3H), 3.57 (t, $J$ = 5.7 Hz, 2H), 2.71 (s, 2H), 1.44 (s, 9H).

**1.6.2. Representative Procedure 5. Preparation of tert-butyl 4-(4-((3-methoxyphenyl)amino)-6-phenylpyrimidin-2-yl)piperidine-1-carboxylate (Ve).**

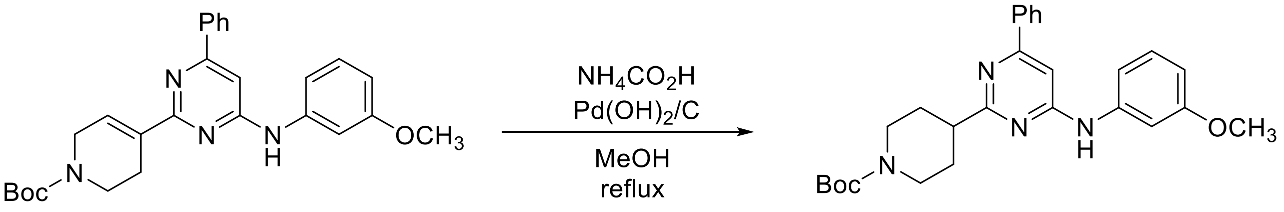


**IVe** **Ve**

Under $N_2$ atmosphere, a suspension of intermediate **IVe** (240 mg, 0.52 mmol, 1 equiv), ammonium formate (131 mg, 2.08 mmol, 4 equiv), $Pd(OH)_2/C$ (48 mg, 20 % of starting material weight) in MeOH dry was stirred at reflux temperature until reaction completion. Catalyst was filtered off through a celite coarse patch and resulting filtrate concentrated to dryness at low pressure. Final normal phase purification (cyclohexane/AcOEt from 100/0 to 80/20) afforded pure title compound **Ve** (120 mg, yield 50 %). Rt = 2.11 min (analysis method 2); MS (ESI) m/z: 461.4 $[M-H]^+$, $[M-H]^+$ calculated: 461.2. $^1H$ NMR (400 MHz, DMSO-$d_6$) δ 9.61 (s, 1H), 8.07 – 7.94 (m, 2H), 7.63 (t, $J$ = 2.2 Hz, 1H), 7.57 – 7.45 (m, 3H), 7.22 (t, $J$ = 8.0 Hz, 1H), 7.17 (dt, $J$ = 8.3, 1.4 Hz, 1H), 7.08 (s, 1H), 6.59 (ddd, $J$ = 8.0, 2.5, 1.1 Hz, 1H), 4.12 – 3.96 (m, 2H), 3.77 (s, 3H), 2.93 (tt, $J$ = 11.4, 3.7 Hz, 3H), 2.06 – 1.95 (m, 2H), 1.72 (qd, $J$ = 12.4, 4.2 Hz, 2H), 1.42 (s, 9H).

**1.6.3. Preparation of *N*-(3-methoxyphenyl)-6-phenyl-2-(piperidin-4-yl)pyrimidin-4-amine (ARN22090)**

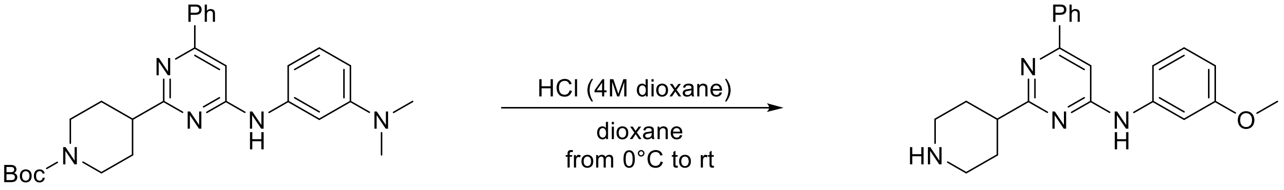


**VIe** **ARN22090**

Compound **ARN22090** was prepared according Representative Procedure 4: using intermediate **VIe**, (115 mg, 0.25 mmol), and HCl (4M) solution in 1,4-dioxane (0.69 ml) in 1,4-dioxane (0.69). Final normal phase purification (elution by gradient from from 95/5 to 50/50 DCM/DCM:$NH_3$ 1M MeOH 4:1) afforded pure title compound **ARN22090** (80.1 mg, yield 89 %). UPLC_MS: Rt = 1.85 min (analysis method 1); MS (ESI) m/z: 361.6 $[M-H]^+$, $[M-H]^+$ calculated: 361.2. $^1H$ NMR (400 MHz, DMSO-$d_6$) δ 9.58 (s, 1H), 8.04 – 8.00 (m, 2H), 7.65 (bs, 1H), 7.55 – 7.47 (m, 3H), 7.25 – 7.19 (m, 2H), 7.07 (s, 1H), 6.58 (dt, $J$ = 7.2, 2.3 Hz, 1H), 3.78 (s, 3H), 3.04 (dt, $J$ = 12.2, 3.3 Hz, 2H), 2.80 (tt, $J$ = 11.9, 3.9 Hz, 1H), 2.61 (td, $J$ = 11.9, 2.5 Hz, 2H), 2.03 – 1.85 (m, 2H), 1.74 (qd, $J$ = 12.2, 4.0 Hz, 2H). $^{13}C$ NMR (100 MHz, DMSO-$d_6$) δ 172.5 (Cq), 161.3 (Cq), 161.2 (Cq), 159.6 (Cq), 141.5 (Cq), 137.3 (Cq),

130.1 (CH), 129.4 (CH), 128.8 (CH, 2C), 126.4 (CH, 2C), 111.6 (CH), 107.6 (CH), 105.0 (CH), 99.7 (CH), 55.0 ($CH_3$), 46.1 ($CH_2$, 2C), 45.6 (CH), 31.8 ($CH_2$, 2C).

#### 1.7. Synthesis of compound ARN22089.

##### 1.7.1. Preparation of tert-butyl 4-(4-((3-(dimethylamino)phenyl)amino)-6-phenylpyrimidin-2-yl)-3,6-dihydropyridine-1(2H)-carboxylate (IVf).

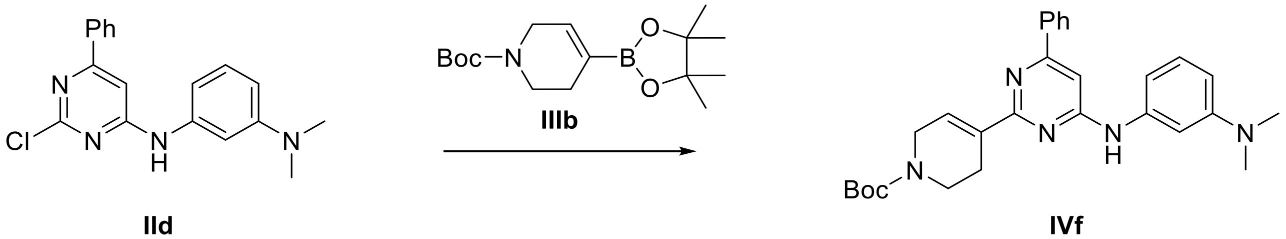


Compound **IVf** was prepared according Representative Procedure 2: using intermediate **IId** (120 mg, 0.37 mmol) and tert-butyl 4-(4,4,5,5-tetramethyl-1,3,2-dioxaborolan-2-yl)-3,6-dihydro-2H-pyridine-1-carboxylate **IIIb** (129.5 mg, 0.41 mmol). Final normal phase purification (cyclohexane/AcOEt from 100/0 to 80/20) afforded title compound **IVf** (155.1 mg, yield 89 %). Rt = 2.48 min (analysis method 2); MS (ESI) m/z 472.3 $[M-H]^+$, $[M-H]^+$ calculated: 472.3. $^1H$ NMR (400 MHz, DMSO-$d_6$) δ 9.44 (s, 1H), 8.17 – 7.95 (m, 2H), 7.67 – 7.41 (m, 3H), 7.31 (s, 1H), 7.21 (s, 1H), 7.14 (t, *J* = 8.1 Hz, 1H), 7.11 (s, 1H), 7.03 – 6.93 (m, 1H), 6.42 (ddd, *J* = 8.3, 2.7, 0.8 Hz, 1H), 4.11 (s, 2H), 3.56 (t, *J* = 5.6 Hz, 2H), 2.93 (s, 6H), 2.81 – 2.62 (m, 2H), 1.44 (s, 9H).

##### 1.7.2. Preparation of tert-butyl 4-(4-((3-(dimethylamino)phenyl)amino)-6-phenylpyrimidin-2-yl)piperidine-1-carboxylate (Vf).

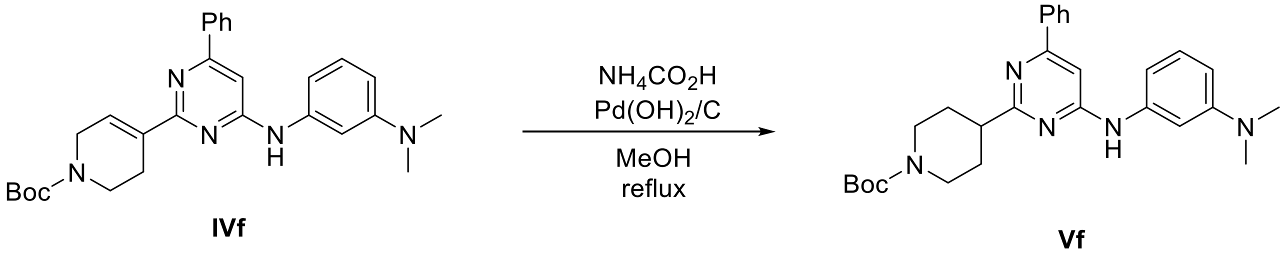


Compound **Vf** was prepared according Representative Procedure 5: using intermediate **IVf** (150 mg, 0.32 mmol). Final normal phase purification (cyclohexane / AcOEt from 100/0 to 80/20) afforded pure title compound **Vf** (149 mg, yield 99 %). Rt = 2.30 min (analysis method 2); MS (ESI) m/z: 474.4 $[M-H]^+$, $[M-H]^+$ calculated: 474.3. $^1H$ NMR (400 MHz, DMSO-$d_6$) δ 9.43 (s, 1H), 8.08 – 7.94 (m, 2H), 7.58 – 7.44 (m, 3H), 7.40 (s, 1H), 7.11 (t, *J* = 8.1 Hz, 1H), 7.06 (s, 1H), 6.88 (ddd, *J* = 7.9, 2.1, 0.8 Hz, 1H), 6.41 (ddd, *J* = 8.4, 2.5, 0.8 Hz, 1H), 4.05 (d, *J* = 13.1 Hz, 2H), 1.98 (dd, *J* = 13.6, 3.5 Hz, 2H), 2.95 – 2.87 (m, 9H), 1.73 (qd, *J* = 12.5, 4.2 Hz, 2H), 1.42 (s, 9H).

##### 1.7.3. Preparation of *N1,N1*-dimethyl-*N3*-(6-phenyl-2-(piperidin-4-yl)pyrimidin-4-yl)benzene-1,3-diamine (ARN22089).

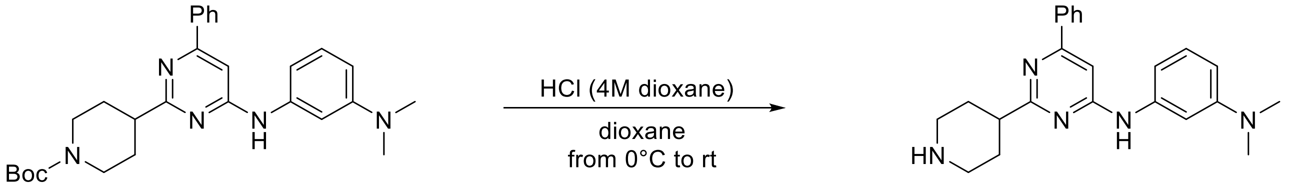


**VIf** **ARN22089**

Compound **ARN22089** was prepared according Representative Procedure 4: using intermediate **VIf**, (80 mg, 0.22 mmol), and HCl (4M) solution in 1,4-dioxane (0.61 ml) in 1,4-dioxane (0.61). Final normal phase purification (elution by gradient from from 95/5 to 60/40 DCM/DCM:$NH_3$ 1M MeOH 4:1) afforded pure title compound **ARN22089** (39.6 mg, yield 34 %). UPLC_MS: Rt = 1.97 min (analysis method 1); MS (ESI) m/z: 374.6 $[M-H]^+$, $[M-H]^+$ calculated: 374.2. $^1$H NMR (400 MHz, DMSO-$d_6$). δ 9.42 (s, 1H), 8.01 (d, $J$ = 7.0 Hz, 2H), 7.51 (d, $J$ = 7.0 Hz, 3H), 7.40 (s, 1H), 7.12 (t, $J$ = 8.1 Hz, 1H), 7.07 (s, 1H), 6.94 (d, $J$ = 8.0 Hz, 1H), 6.51 – 6.33 (m, 1H), 3.03 (d, $J$ = 11.9 Hz, 2H), 2.93 (s, 6H), 2.85 – 2.70 (m, 1H), 2.61 (t, $J$ = 11.9 Hz, 2H), 2.00 – 1.85 (m, 2H), 1.75 (qd, $J$ = 12.3, 4.3 Hz, 2H). $^{13}$C NMR (100 MHz, DMSO-$d_6$) δ 161.9 (Cq), 161.5 (Cq), 151.4 (Cq), 141.4 (Cq), 137.9 (Cq), 130.5 (CH), 129.5 (CH), 129.2 (CH, 2C), 126.4 (CH, 2C), 108.3 (CH), 107.2 (CH), 104.3 (CH), 100.0 (CH), 45.8 ($CH_2$, 2C), 45.2 (CH), 40.2 ($CH_3$), 31.4 ($CH_2$, 2C).

**1.7 Binding evaluation of hit and derivative to CDC42 by MST.**

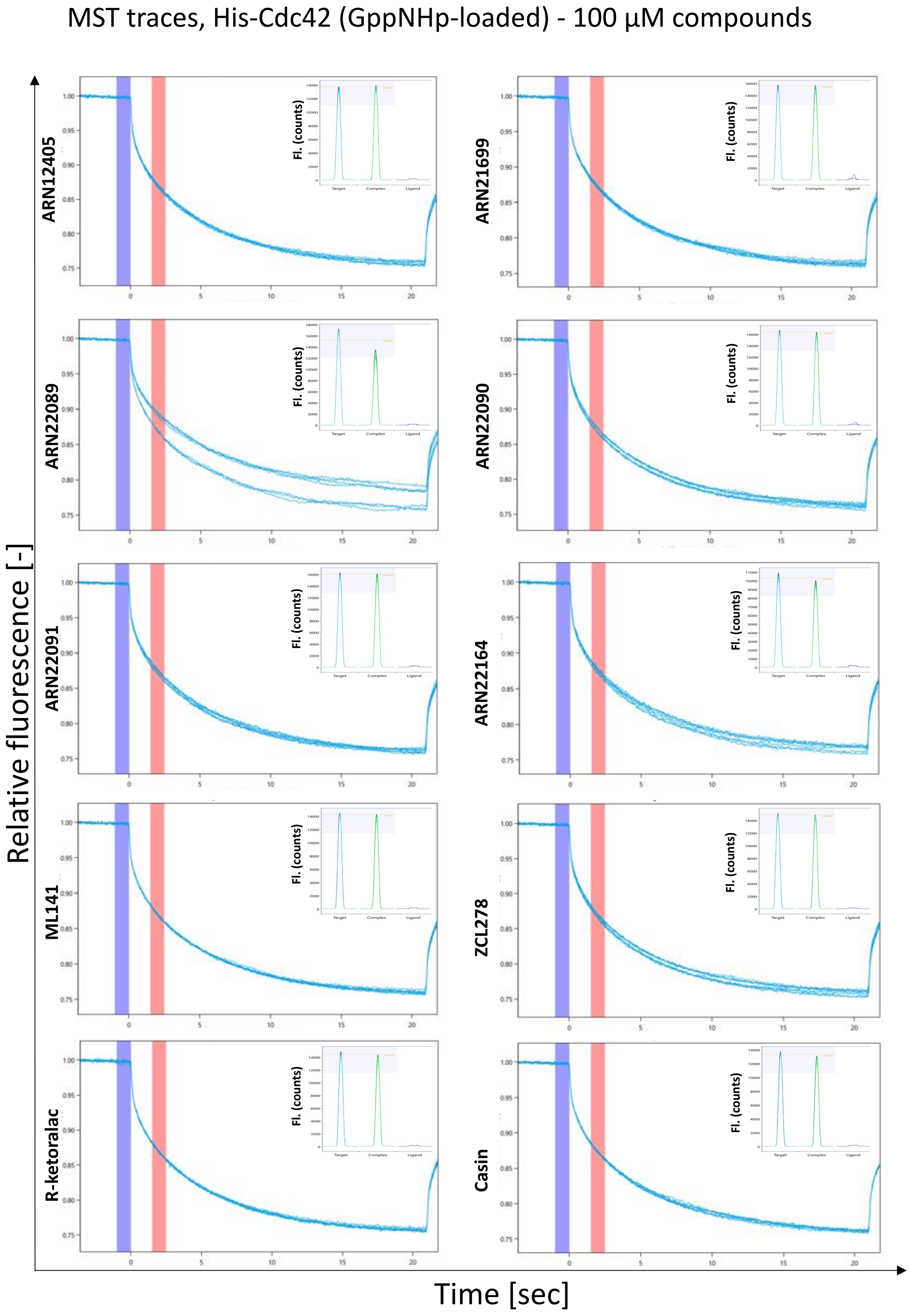

MST traces, His-Cdc42 (GDP-loaded) - 100 µM compounds

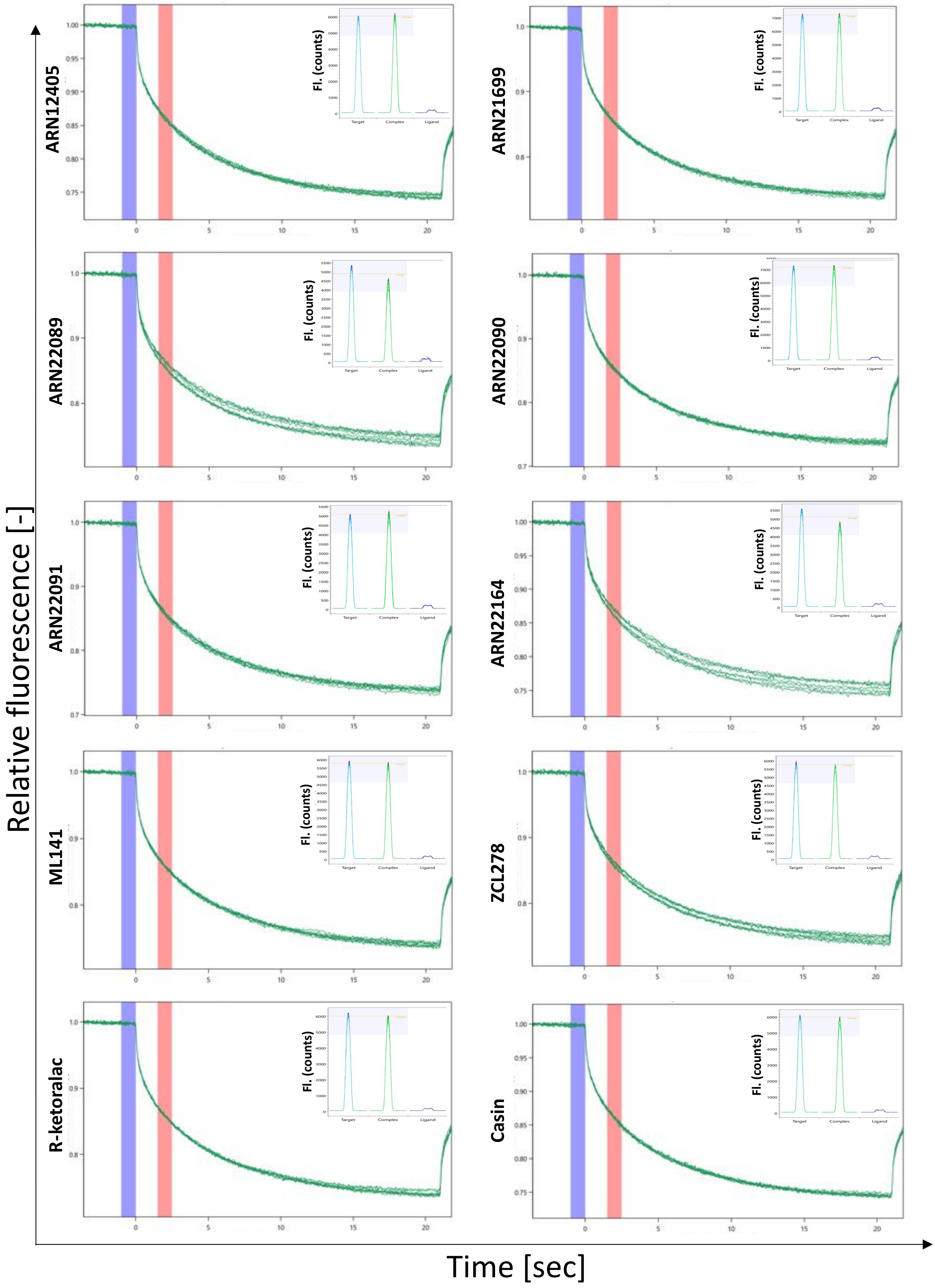

MST traces, His-Cdc42 (GppNHp-loaded) - 50 µM compounds

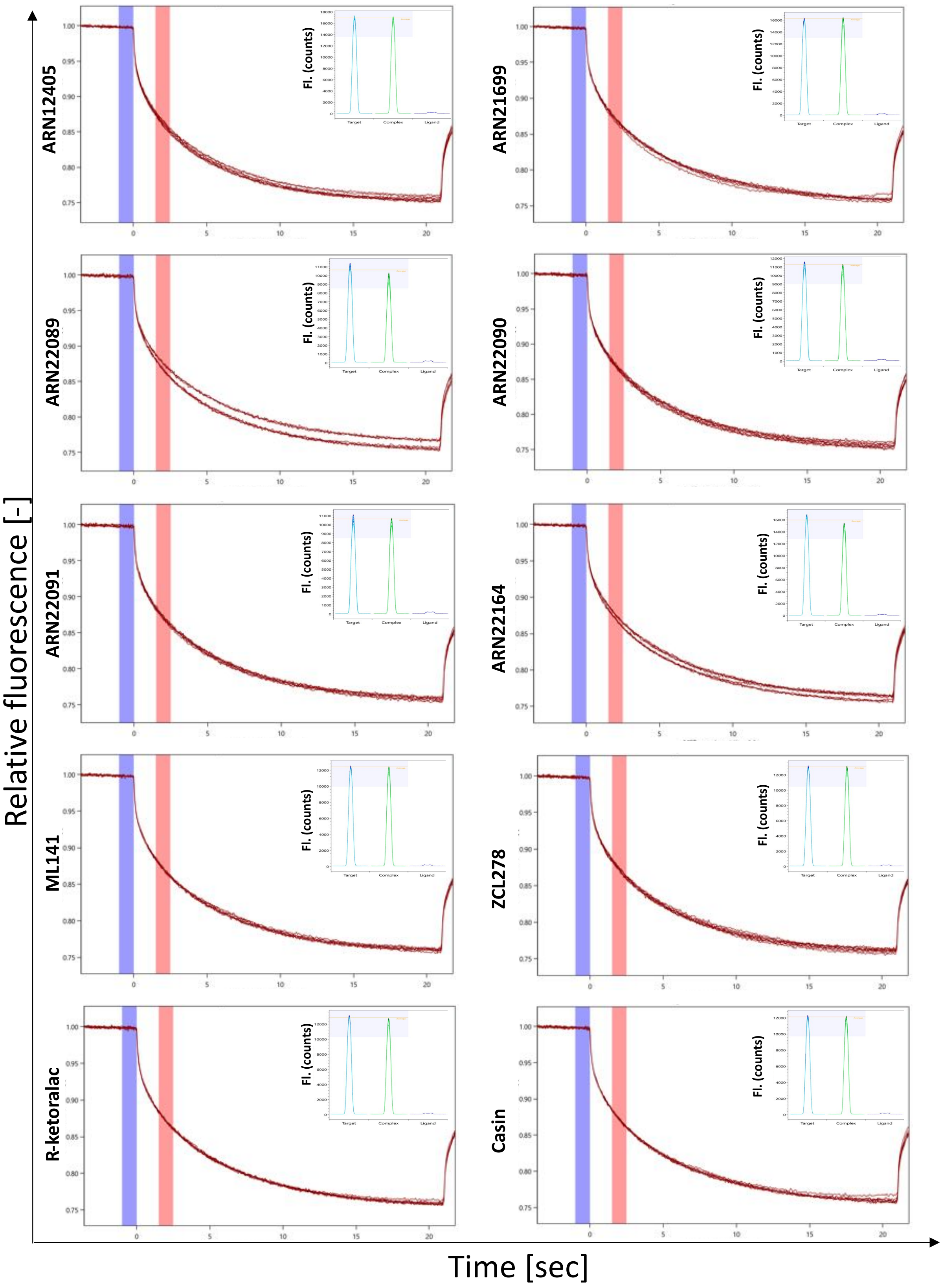

MST traces, His-Cdc42 (GDP-loaded) - 50 μM compounds

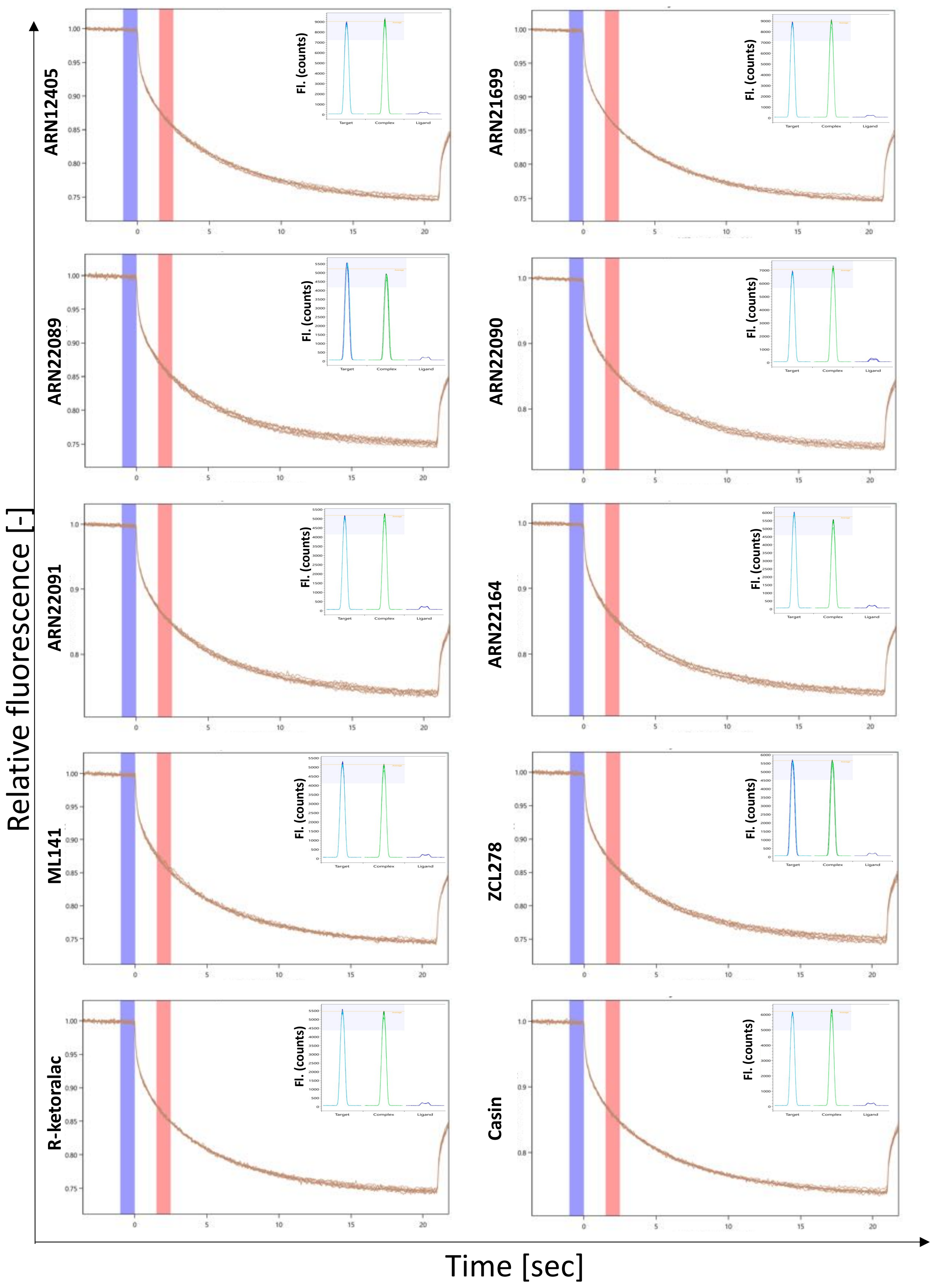

Thermographs of Cdc42 in Trizma pH 7.5, 40 mM NaCl, 0.05% v/v Tween20 buffer, 0.5 or 1 % v/v DMSO. Significant aggregation was not detected after 20 sec of IR laser heating at 20 % MST power. Cold region is set to 0 sec (blue) and hot region to 2.5 sec (pink). Fluorescence counts of target-only, complex and ligand-only samples are also reported in the top right corner of each graph.

**1.8 Evaluation of protein:ARN22089 complex fluorescence change by SD-TEST.**

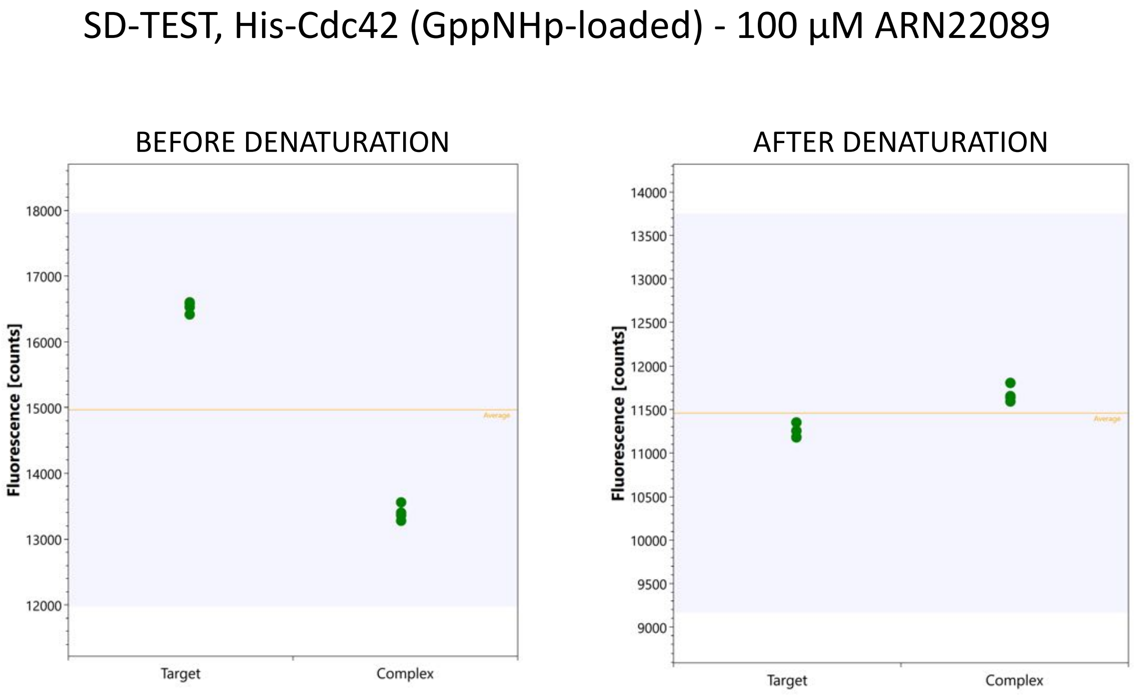


Folded and denatured target-only and complex samples are compared. For denaturation, samples were diluted 1:1 with Tris pH 7.5, 4 % v/v SDS, 40 mM DTT and boiled at 95 °C for 15 min. 11.3 % fluorescence change was detected before denaturation, while 3 % fluorescence change was detected after denaturation. The ligand induced fluorescence change is binding-specific.
